# Optimizing a Culture-Enriched Hybrid Metagenomics Pipeline to Assess the AMR Footprint of Livestock Manure in Anaerobic Digestate

**DOI:** 10.64898/2026.04.24.720626

**Authors:** Nahidur Rahman, A S M Zisanur Rahman, David B. Levin, Tim A. McAllister, Nazim Cicek, Hooman Derakhshani

**Author notes:** Nahidur Rahman and A. S. M. Zisanur Rahman contributed equally to this work.

## Abstract

The role of environmental samples from livestock production systems, including manure and anaerobic digestate, as reservoirs of antimicrobial resistance genes (ARGs) is likely underestimated because conventional metagenomic approaches can overlook low-abundance ARGs and often lack the resolution needed to reliably associate these genes with their microbial hosts and linked mobile genetic elements (MGEs). Here, we evaluated whether culture-enriched metagenomics (CEMG), with and without antibiotic selection, enhances ARG detection in anaerobic digestate and improves the resolution of ARG-MGE-host associations using hybrid short- and long-read metagenomic assembly. Culture enrichment substantially increased ARG recovery, mean ARG abundance rose from 15.4 counts per million (CPM) in culture-independent direct metagenomes from fresh digestate (FD) to 124 CPM in CEMG without antibiotics and 160.0 CPM in antibiotic-selective CEMG, corresponding to an approximately 10.4-fold increase over FD. In FD, only 9 unique ARGs were detected, whereas enrichment recovered 112, including ARGs of clinical importance such as glycopeptide resistance, β-lactamase genes of the CTX-M, OXA, and TEM families, and the *cfr* 23S rRNA methyltransferase conferring cross- resistance to multiple antibiotic classes. Oxygen availability was the strongest factor structuring enriched community compositions and ARG profiles, with aerobic and anaerobic communities forming distinct clusters. Antibiotic selection induced targeted, class-specific shifts in ARG profiles, with ARGs associated with tetracycline resistance consistently enriched across treatments. Hybrid metagenomic assembly resolved the genomic context of 784 ARGs, of which 59.3% were co-localized with at least one class of mobile genetic element (MGE), predominantly plasmids, insertion sequences, and integrative and conjugative/mobilizable elements (ICEs/IMEs). Biocide and metal resistance genes frequently co-occurred with ARGs on the same contigs, highlighting the potential for co-selection. Together, these findings demonstrate that antibiotic-selective culture enrichment enhances resistome surveillance by improving detection of low-abundance ARGs, while hybrid assembly provides critical genomic context for assessing their mobility and host associations.

**IMPORTANCE:** Livestock manure and its byproducts, such as anaerobic digestate, are recognized as important environmental reservoirs of antimicrobial resistance genes and resistant bacteria, yet current metagenomic approaches may underestimate this risk by failing to detect low abundance but clinically relevant ARGs. Here, we show that integrating culture enrichment with hybrid metagenomics improves ARG recovery and reveals ARG co-localization with mobile genetic elements and putative bacterial hosts. This approach captures a cultivable and condition-responsive fraction of the resistome that is not readily accessible through direct metagenomic sequencing alone, providing a more informative framework for environmental AMR surveillance.

## INTRODUCTION

Antimicrobials are increasingly recognized as emerging environmental contaminants in diverse ecosystems, including soil and aquatic environments, because of their role in driving the emergence and dissemination of antimicrobial resistance (AMR) (1). A substantial proportion of these contaminants originates from human and veterinary antimicrobial use (2). On average, over 100 mg of antimicrobials were used globally to produce each kilogram of animal biomass in 2016 (3). Notably, some of the antimicrobial classes used in animals, including macrolides, fluoroquinolones, and cephalosporins, are designated as critically important for treating infectious diseases in humans (4). Antimicrobial use in food-producing animals selects for resistant bacteria and antimicrobial resistance genes (ARGs) within the gastrointestinal tract. Because many antimicrobials are only partially metabolized or absorbed in the gut, a substantial proportion is excreted in urine and feces alongside resistant bacteria and ARGs. Consequently, manure serves as a reservoir of resistant bacteria, ARGs, and antimicrobial residues that can further select and co-select for AMR in environmental microbial communities (5). Digestate, the byproduct of anaerobic digestion of manure, is widely used as a fertilizer in integrated livestock and cropping systems. Although anaerobic digestion reduces pathogen abundance and the overall load of antimicrobial resistance genes, the resulting digestate may still retain residual resistance genes and associated elements that can be disseminated into the environment following land application (5, 6). Characterizing the resistome of these reservoirs requires sensitive, high throughput approaches capable of detecting diverse resistance determinants. Direct culture-independent metagenomics, which enables sequencing of environmental DNA without the need for cultivation (7), has become a widely adopted strategy, providing broader and more comprehensive ARG profiling in complex environmental samples than targeted approaches such as high-throughput quantitative PCR (8–11). It also enables the simultaneous characterization of microbial diversity, resistome profiles, and mobile genetic elements (MGEs) such as plasmids, transposons, and phages (12).

Despite this practical advantage, direct metagenomics has important limitations for resistome surveillance. Its effectiveness is constrained by the sequencing platform, community complexity, and reference-database quality, and it frequently fails to detect underrepresented or low-abundance microbial taxa (13, 14), yielding incomplete representations of ARGs and MGEs in complex environments such as wastewater, manure, and digestate (15, 16). This blind spot is consequential because low-abundance microbial taxa can act as reservoirs of clinically relevant ARGs (17), which can be mobilized and transferred to abundant or potentially pathogenic community members through horizontal gene transfer (18). Moreover, these ARG-carrying taxa may expand when conditions change, for example, following land application of manure or digestate. Methods that systematically under-detect rare populations therefore risk underestimating both the diversity and the dissemination potential of the resistome. Recovering this rare fraction by direct metagenomics typically demands ultra-high sequencing depth at substantial cost (19). Hybridization-based bait capture can enrich low-abundance ARGs, but it introduces selection biases and depends on careful probe design and accurate on- and off-target discrimination. In addition, when combined with short-read sequencing, it still limits recovery of the genomic context needed to resolve ARG-MGE associations (20).

Selective culture enrichment integrated with direct metagenomics provides a complementary strategy that extends the scope of resistome characterization. Rather than replacing direct metagenomics, this approach targets the cultivable fraction of the resistome, enhancing the detection and recovery of rare, resistant populations that may be underrepresented or overlooked in culture-independent analyses. This strategy has enhanced resistome characterization across environments ranging from the human gut (17) and cystic fibrosis sputum (9) to drinking water (21), wastewater (19) and ocean sediments (22). By favoring the growth of targeted groups such as antimicrobial-resistant bacteria (19) and capturing previously uncultured or poorly characterized microbes (22, 23), enrichment increases the sensitivity of downstream analyses (14), a particular advantage in AMR research, where ARGs constitute less than 1% of bacterial DNA (10) and only about 0.05% of metagenomic reads (24).

Improved ARG detection alone, however, does not resolve the critical question of which organisms carry these determinants. Illumina short-read sequencing has limited ability to link ARGs to their microbial hosts and can be affected by PCR amplification bias (25). Moreover, short-read assembly into contigs can further compromise ARG recovery, as assemblies often fragment across repetitive and conserved regions flanking ARGs (18) and fail to resolve the complex, mosaic architectures of MGEs (26, 27), the very genetic structures that determine the potential for ARG mobility and dissemination (18). Third-generation long-read platforms, such as Oxford Nanopore and PacBio, generate reads spanning tens of thousands of bases that can cover full-length ARGs and their flanking regions, aiding host inference and MGE resolution (25). Although their per-base accuracy has improved substantially with recent chemistries, combining long-read and short-read data remains advantageous, as the high base-level accuracy and depth of short reads complement the contiguity of long reads (26). Integrating this hybrid sequencing strategy with culture enrichment can further enhance host identification and resolve the genomic context of ARGs, particularly those embedded within or associated with MGEs.

Digestate is an anoxic environment dominated by obligate and facultative anaerobes (28, 29), so recovering its cultivable resistome requires anaerobic cultivation, while parallel aerobic enrichment makes it possible to assess how oxygen availability shapes which resistance determinants are recovered. Building on this rationale, our study had two objectives: (i) to compare direct culture-independent metagenomics and culture-enriched metagenomics (CEMG) for their ability to characterize the resistome and microbial community structure of anaerobic digestate, while optimizing enrichment conditions across oxygen regimes, antimicrobial classes, and concentrations to enhance ARG recovery; and (ii) to integrate hybrid short- and long-read assembly into this workflow to resolve the genomic context of ARGs, including their associations with MGEs and bacterial hosts.

## RESULTS

Across 81 samples comprising fresh digestate collected at three time points (FD, n = 3), non-selective culture-enriched metagenomes (CEMG-NoAB, n = 6), and antibiotic-selective culture-enriched metagenomes (CEMG-AB, n = 72) (Table S1), Illumina sequencing yielded approximately 1.94 billion reads (968 million read pairs). FD libraries yielded an average of 63.8 ± 2.4 million reads per sample, whereas CEMG-NoAB and CEMG-AB yielded 19.2 ± 9.3 and 22.6 ± 7.5 million reads per sample, respectively (mean ± SD; Table S2). Culture-enriched libraries were sequenced to lower depth than direct metagenomes. We analyzed these data using two complementary approaches: read-based mapping to reference databases, to profile microbial community and ARG composition, and hybrid short- and long-read assembly, to resolve the genomic context of ARGs and their associations with MGEs and microbial hosts.

### The impact of culture enrichment on microbial diversity of digestate

To provide biological context for the downstream resistome analyses, we first assessed how culture enrichment altered the taxonomic composition of the FD microbial community. At the genus level, the three FD replicates - collected from the same digester at different time points - clustered tightly, reflecting a stable, homogeneous baseline community (Fig. 1A). Culture-enriched communities (CEMG-NoAB and CEMG-AB) diverged markedly from FD (PERMANOVA, R^2^ = 0.068, *p* = 0.001).

**Figure 1.**
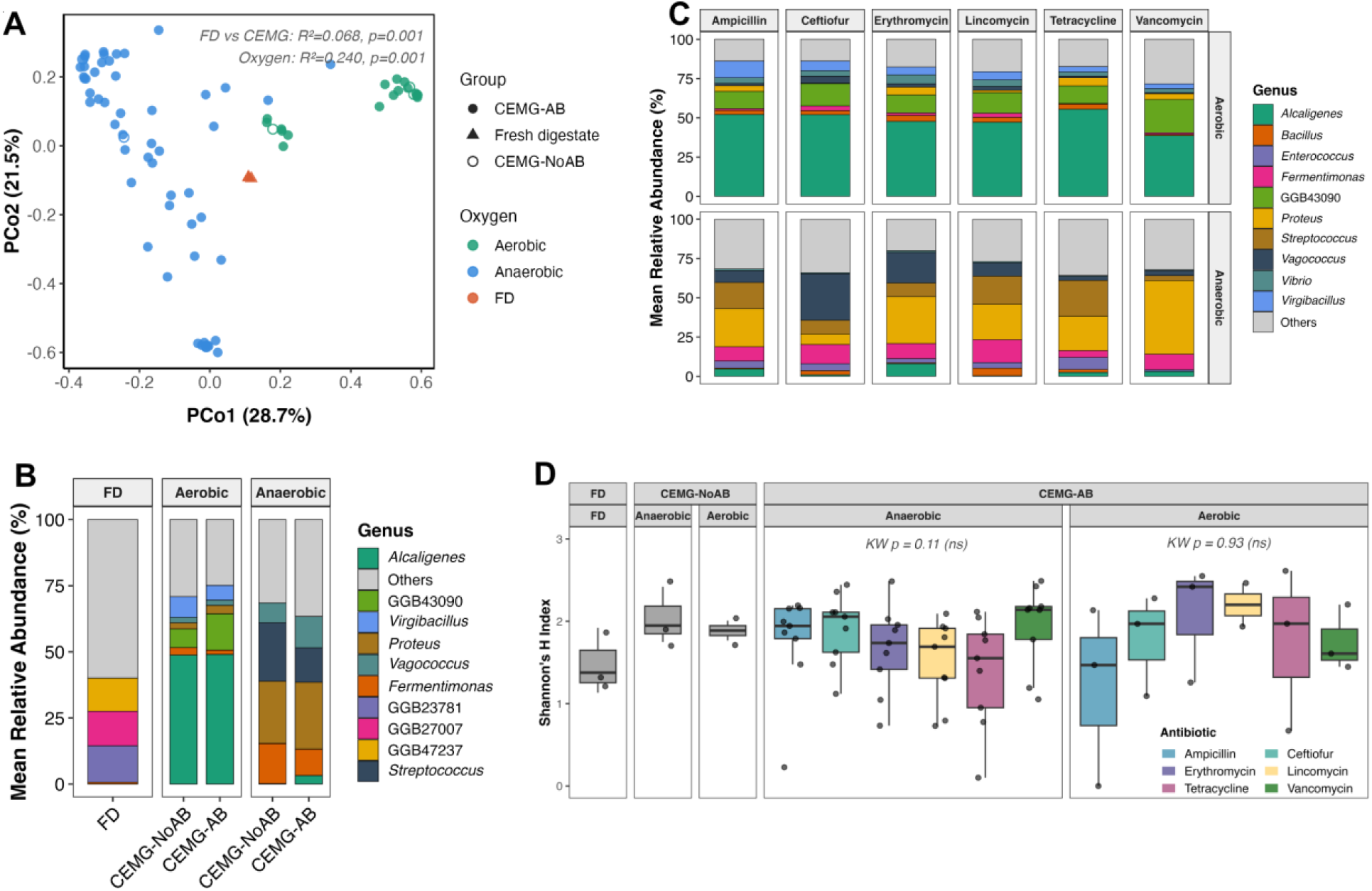
Taxonomic restructuring of digestate microbial communities following culture enrichment. **(A)** Principal coordinates analysis (PCoA) of Bray-Curtis dissimilarity at the genus level. Points are coloured by oxygen condition (aerobic, teal; anaerobic, blue; FD, orange) and shaped by sample group (filled circle: CEMG-AB; open circle: CEMG-NoAB; triangle: fresh digestate). PERMANOVA results are annotated within the panel. **(B)** Mean relative abundance of the top ten genera across FD, aerobic (CEMG-NoAB and CEMG-AB), and anaerobic (CEMG-NoAB and CEMG-AB) communities; remaining genera are grouped as “Others”. GGB entries denote unclassified genome-based genus-level bins. **(C)** Mean relative abundance of the top genera across the six antibiotic enrichment treatments (ampicillin, ceftiofur, erythromycin, lincomycin, tetracycline, vancomycin), stratified by oxygen condition (aerobic, upper panels; anaerobic, lower panels). **(D)** Shannon diversity index across FD, CEMG-NoAB, and CEMG- AB, stratified by oxygen condition and antibiotic treatment. Data points represent individual samples; boxplots summarize within-group distributions. Differences were assessed using the Kruskal-Wallis test, followed by Dunn’s post hoc test with Benjamini-Hochberg correction for multiple comparisons.

Oxygen exposure was the strongest driver of enriched-community composition, with aerobic and anaerobic communities showing clear clustering patterns in principal coordinates analysis (PERMANOVA, R^2^ = 0.240, *p* = 0.001; Fig. 1A). Aerobic communities exhibited a trend toward lower within-group dispersion compared with anaerobic communities, although this difference was not statistically significant (PERMDISP, *p* = 0.069; mean distance to centroid: aerobic = 0.40, anaerobic = 0.48; Fig. S1A). This pattern was consistent with aerobic enrichment converging toward a narrow, *Alcaligenes*-dominated community profile across antibiotic treatments, whereas anaerobic communities displayed greater compositional variability among enrichment conditions (Fig. S1B). Aerobic communities were consistently dominated by *Alcaligenes* (∼50.2%), with contributions from an unclassified member of *Alcaligenaceae* (GGB43090; ∼12.3%), *Virgibacillus* (∼6%), *Halopseudomonas* (∼4%), *Vibrio* (∼4%), and *Proteus* (∼3%). Anaerobic communities were enriched in *Proteus* (∼25%), *Streptococcus* (∼13%), *Vagococcus* (∼11%), and *Fermentimonas* (∼10.5%) (Fig. S1B).

FD displayed a distinct taxonomic profile dominated by unclassified genome-based genus-level bins (GGBs) affiliated primarily with the phyla *Bacteroidota* (∼41%) and *Mycoplasmatota* (∼26%) (Fig. 1B). Within culture-enriched groups, community composition was broadly consistent between CEMG-NoAB and CEMG-AB under the same oxygen regime, with aerobic communities dominated by *Alcaligenes* and anaerobic communities showing *Proteus*-, *Streptococcus*-, and *Fermentimonas*-dominated profiles.

Individual antibiotics produced minor differences in dominant taxonomic abundances, with the same primary genera consistently observed under each oxygen condition (Fig. 1C). Community diversity (Shannon index) did not differ significantly among the six antibiotic treatments under either anaerobic (Kruskal-Wallis, *p* = 0.11) or aerobic (*p* = 0.93) enrichment conditions (Fig. 1D) or across antibiotic concentration under anaerobic conditions (Fig. S1C). However, both enriched groups showed significantly lower Shannon diversity than FD (mean: FD = 3.02, CEMG-NoAB = 1.63, CEMG-AB = 1.68; Kruskal-Wallis, *p* = 0.014), with post-hoc Dunn tests (Benjamini-Hochberg correction) confirming that FD differed significantly from both CEMG-NoAB (*p* = 0.020) and CEMG-AB (*p* = 0.011), while CEMG-NoAB and CEMG-AB did not differ from each other (*p* = 0.93; Fig. 1D). Among the antibiotic concentrations evaluated for dose-response enrichment, only vancomycin exhibited a significant concentration-dependent effect, with the number of recovered genera declining progressively as antibiotic concentration increased (linear fit, *p* = 0.003, R^2^ = 0.74; model selection described in Methods; Table S4). Overall, the lowest tested concentration (1X) preserved number of unique genera detected per sample across antibiotics, with ceftiofur as the sole exception, where richness peaked at 10X before declining.

CEMG-AB expanded detectable species-level diversity. A total of 145 species were identified exclusively in CEMG-AB samples and were absent from the direct metagenomics of FD as well as CEMG-NoAB. Among these, 91 species were detected in at least two independent CEMG-AB samples, providing a conservative estimate of taxa specifically enriched by antibiotic selection while minimizing potential single-sample artifacts (Table S5). This fraction included clinically and environmentally relevant species such as *Enterococcus dispar*, *Enterobacter hormaechei*, *Citrobacter amalonaticus*, *Acinetobacter towneri*, *Enterococcus casseliflavus*, *Enterococcus faecalis*, and *Achromobacter insolitus*, among others.

### CEMG-derived changes in resistome composition and diversity

Direct alignment of quality-controlled Illumina reads to the CARD database revealed a markedly higher ARG abundance in CEMG-AB relative to FD (Table S6). Mean ARG abundance increased from 15.4 CPM (median 14.1) in FD to 160.0 CPM (median 139.0) in CEMG-AB, corresponding to a 10.4-fold increase despite lower sequencing depth in enriched libraries. CEMG-NoAB also showed substantially higher ARG abundance (mean 124 CPM) relative to FD, suggesting that CEMG, even in the absence of antibiotic selection, can recover a cultivable resistome that is poorly represented in direct metagenomics (Table S6).

Presence-absence analysis further demonstrated the enhanced ARG detection capacity of CEMG. Across all samples, 116 unique high-confidence ARGs were identified based on stringent criteria (>98% nucleotide identity and ≥90% coverage against the CARD database). Of these, only 9 ARGs were detected in direct metagenomics of FD, including 5 that were shared with CEMG (Fig. 2). In contrast, CEMG recovered 112 ARGs, accounting for approximately 96.6% of all detected ARGs. Notably, several clinically relevant ARG families were detected exclusively in enriched samples. These included members of the *VanH, VanS, VanT, VanXY*, and Van ligase gene families, which are suggestive of glycopeptide resistance potential; β-lactamase genes of the CTX-M, OXA, and TEM families, variants of which can confer extended-spectrum β- lactamase (ESBL) activity; the multi-class rRNA methyltransferase gene *cfr*, which can confer resistance to phenicols, lincosamides, oxazolidinones, pleuromutilins, and streptogramin; Isa- type ABC-F proteins, which can mediate pleuromutilin, lincosamide, and streptogramin resistance; and multiple aminoglycoside-modifying enzyme genes (*ant*(3’), *ant*(4’), *ant*(9), *aac*(6’), *aph*(2’)).

**Figure 2.**
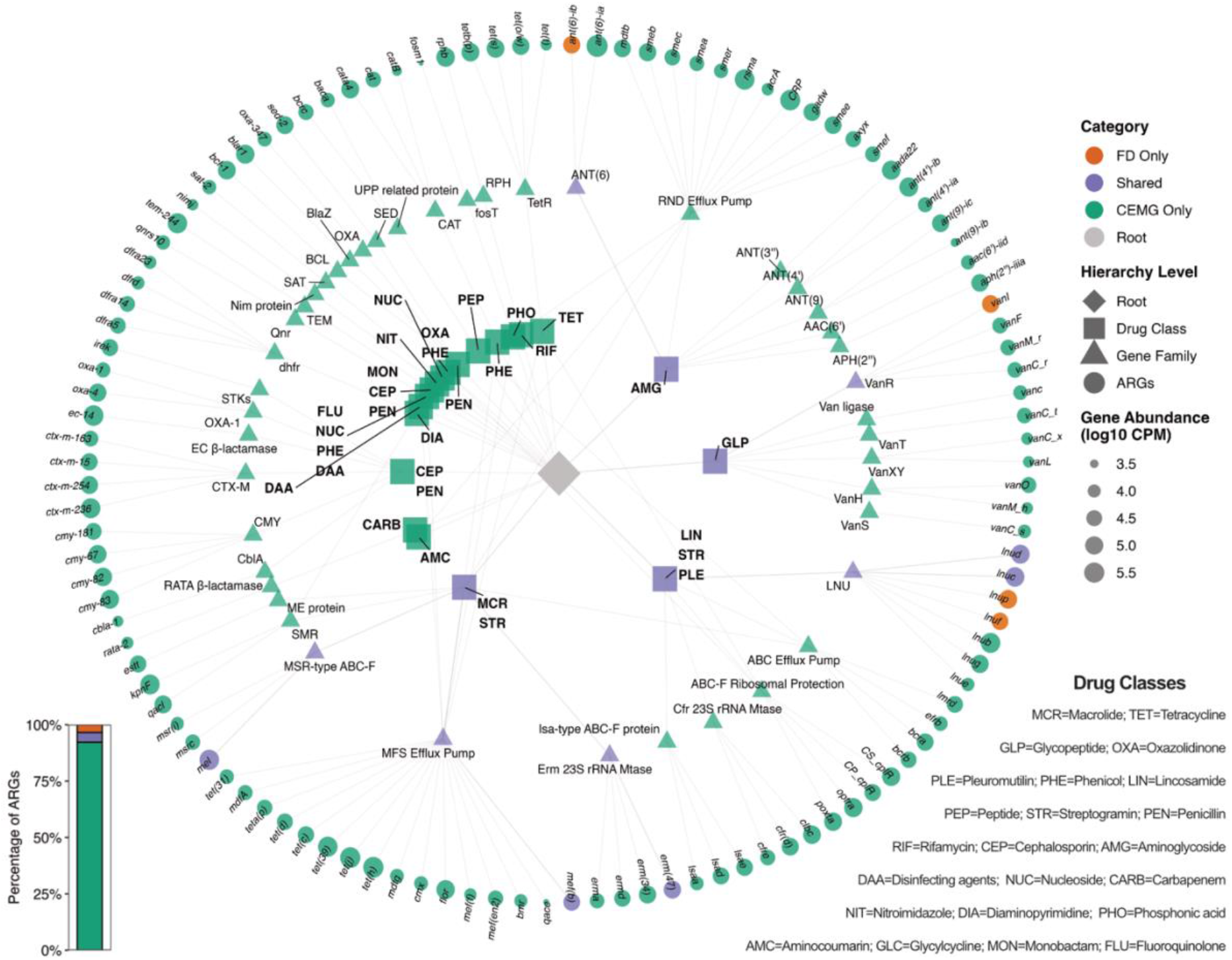
Network representation of antimicrobial resistance genes (ARGs) detected across treatment groups. ARGs are visualized as a hierarchical network read from center outward: the root node (diamond) connects to resistance drug classes (squares), which link to ARG gene families (triangles), which in turn connect to individual ARG genes (circles) in the outer ring. Node colors indicate group-specific occurrence based on presence-absence across samples: orange = FD only; purple = shared between FD and CEMG; teal = CEMG only. Node sizes reflect mean ARG abundance (log_10_ counts per million reads; CPM). The inset bar plot summarizes the number of ARGs in each occurrence category (FD only, shared, CEMG only) as a proportion of all detected ARGs.

The richness of ARGs associated with clinically important drug classes was significantly lower in FD than in anaerobic CEMG-AB (Fig. 3A). Strong shifts in dominant ARG drug class profiles were observed among FD, CEMG-NoAB, and CEMG-AB (Fig. 3B). The FD resistome was dominated by ARGs associated with macrolide-lincosamide-streptogramin (MLS) resistance, collectively accounting for 92.9% of total ARG-mapping reads, with genes associated with lincosamide resistance (44.1%), macrolide-streptogramin resistance (39.6%), and the combined MLS class (9.2%) together comprising the near-entirety of the FD ARG complement. This composition was consistent with the high relative abundance of *mel* and *erm*(47) genes in FD (Fig. 3C). In contrast, CEMG samples displayed a broader ARG drug-class profile. Under anaerobic CEMG-NoAB conditions, ARGs associated with tetracycline resistance were most prominent (39.4%), followed by ARGs associated with macrolide-streptogramin (8.1%), phenicol (7.9%), lincosamide (6.8%), and oxazolidinone-phenicol (6.5%) resistance (Fig. 3B). Under aerobic CEMG-NoAB conditions, ARGs associated with penicillin resistance were most abundant (26.2%), followed by macrolide-streptogramin (24.6%), MLS (11.2%), and aminoglycoside (10.6%) resistance classes (Fig. 3B). CEMG-AB was dominated by ARGs associated with tetracycline resistance under both aerobic (24.6%) and anaerobic (39.8%) conditions (Fig. 3B). Under aerobic CEMG-AB, aminoglycoside (15.3%) and penicillin (14.4%) resistance ARGs were the next most abundant classes. Under anaerobic CEMG-AB, oxazolidinone-phenicol (8.9%) and macrolide-streptogramin (7.8%) ARGs were the next most prominent, alongside a substantial unclassified fraction (Others: 16.9%) (Fig. 3B). This compositional shift was accompanied by increased abundances of multiple ARGs including *tet*(J), *tet*(H), *mel*, *ant*(4’)-Ib, *cplR*, and *rphB*, under both oxygen conditions (Fig. 3C).

**Figure 3.**
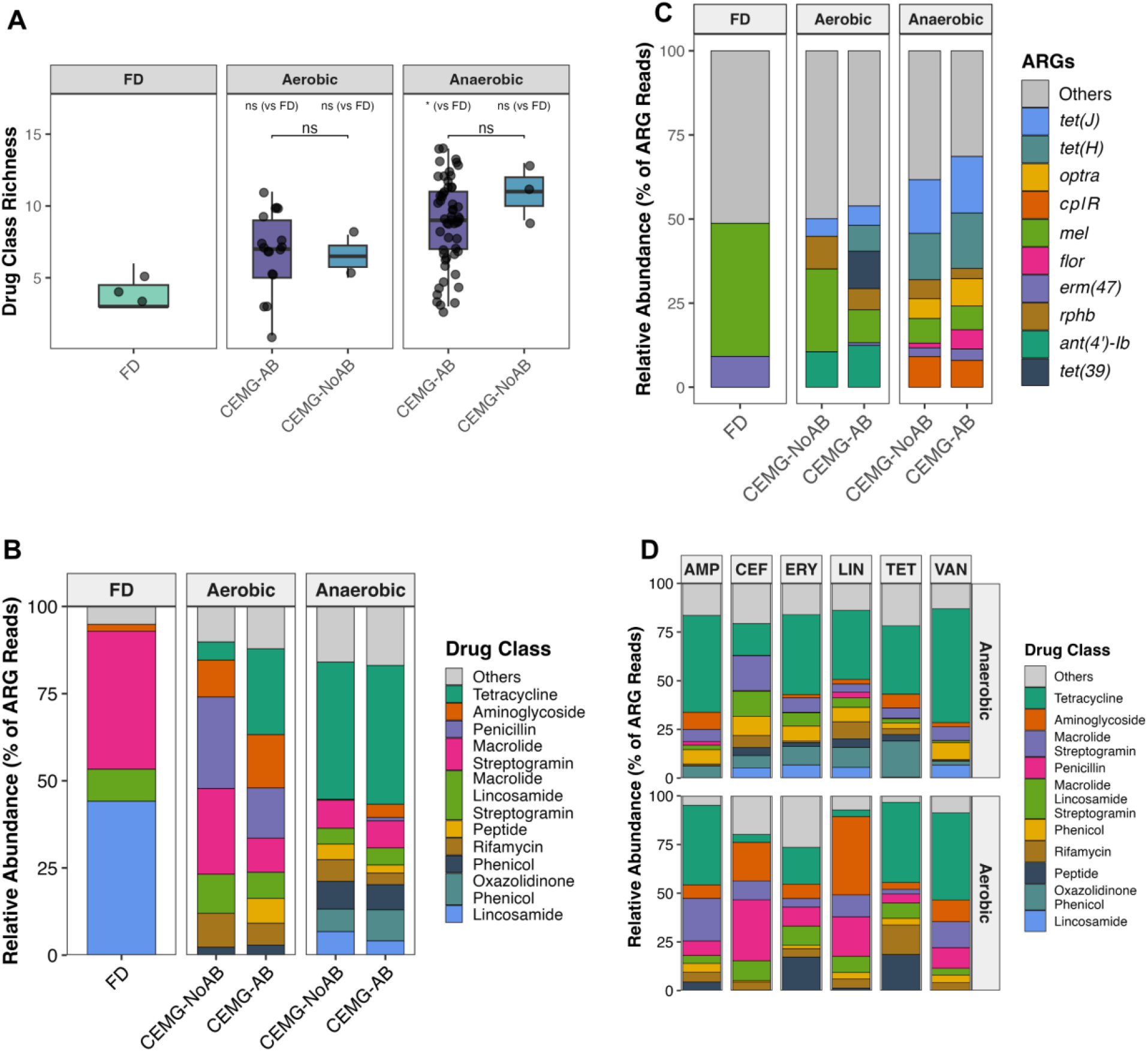
ARG diversity and drug-class composition across enrichment conditions and antibiotic treatments. **(A)** ARG drug class richness under aerobic and anaerobic conditions across FD, CEMG-NoAB, and CEMG-AB. Data points represent individual samples; boxplots summarize within-group distributions. Pairwise group differences were assessed using the Wilcoxon rank-sum test; significance levels are indicated as ns (not significant) and *p* < 0.05 (*). **(B)** Mean relative abundance of the top 10 ARG drug classes across FD, CEMG-NoAB, and CEMG-AB, expressed as a percentage of total ARG-mapping reads within each group; remaining classes are grouped as “Others”. Drug class labels follow CARD annotations. ARGs assigned to multiple drug classes by CARD were retained according to their individual resistance classifications and were not collapsed into a single category. **(C)** Mean relative abundance of the top 10 ARG genes across FD, CEMG-NoAB, and CEMG-AB, expressed as a percentage of total ARG-mapping reads within each group; remaining genes are grouped as “Others”. **(D)** Mean relative abundance of ARG drug classes across antibiotic enrichment treatments (AMP = Ampicillin; CEF = Ceftiofur; ERY = Erythromycin; LIN = Lincomycin; TET = Tetracycline; VAN = Vancomycin), expressed as a percentage of total ARG-mapping reads within each treatment group, shown separately for anaerobic (upper) and aerobic (lower) conditions.

Across different antibiotic-enrichment treatments, ARG profiles were dominated by a small set of recurrent drug class categories, though the relative dominance of specific classes varied with both oxygen condition and antibiotic treatment (Fig. 3D). Under anaerobic conditions, ARGs associated with tetracycline resistance were the most abundant class across all six antibiotic treatments, ranging from 16.3% (ceftiofur) to 58.4% (vancomycin) of total ARG-mapping reads. Under aerobic conditions, tetracycline resistance ARGs were similarly dominant for ampicillin (40.9%), tetracycline (41.1%), and vancomycin (44.7%) enrichments, but were markedly less prominent under ceftiofur (4.1%) and lincomycin (3.3%) enrichments, where penicillin (31.3% and 20.4%, respectively) and aminoglycoside (19.9% and 40.2%, respectively) resistance ARGs instead predominated. Despite this treatment-specific variation, ARGs associated with tetracycline resistance were the most consistently enriched class overall, ranking as the dominant class across all six anaerobic and three of six aerobic antibiotic treatments (AMP: 40.9%, TET: 41.1%, VAN: 44.7%) (Fig. 3D).

Ordination of ARG profiles demonstrated clear compositional differences between aerobic and anaerobic enrichment conditions (Fig. S3A), indicating that oxygen availability strongly shaped resistome composition during culture enrichment. In contrast, ARG profiles among the six antibiotic enrichment treatments did not cluster separately (Fig. S3B), reflecting broadly similar resistome structures across antibiotics. Nevertheless, pairwise PERMANOVA revealed selective compositional differences between specific antibiotic treatments, particularly tetracycline versus vancomycin (*p* = 0.049) and ceftiofur relative to vancomycin (*p* = 0.005) and tetracycline (*p* = 0.014).

Consistent with the taxonomic diversity patterns observed under different antibiotic concentrations, only vancomycin showed a significant dose-dependent association with ARG richness, displaying a negative linear trend (slope = −0.076, p = 0.034; R^2^ = 0.50). Ampicillin, ceftiofur, tetracycline, erythromycin, and lincomycin did not exhibit significant dose-dependent trends (all *p* > 0.1; Fig. S4).

Overall, ARGs conferring tetracycline resistance showed the highest increase in abundance (∼14- fold) relative to FD, with lincomycin enrichment conditions selecting most strongly for communities enriched in tetracycline resistance ARGs (Fig. 4). Similarly, ARGs associated with cephalosporin resistance increased approximately 13-fold relative to FD, with vancomycin and lincomycin enrichment showing the strongest selective effect. Vancomycin enrichment was also associated with the largest increases in aminoglycoside and β-lactam resistance ARGs. ARGs associated with phenicol resistance increased approximately 6-fold following enrichment, with ampicillin and lincomycin conditions contributing most to this increase. Glycopeptide resistance ARGs increased approximately 5.5-fold, while fluoroquinolone and diaminopyrimidine resistance ARGs each increased approximately 5-fold relative to FD.

**Figure 4.**
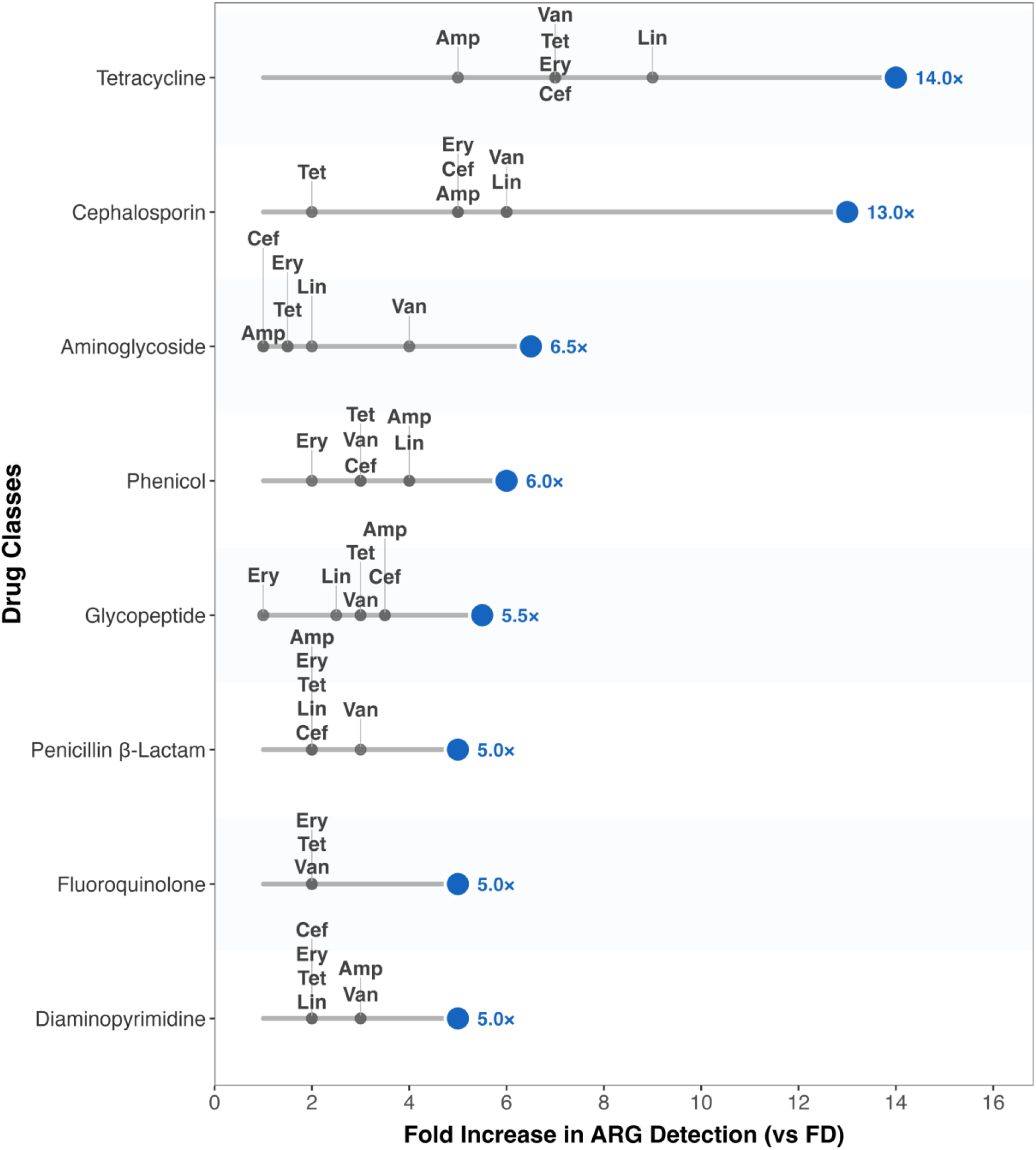
Fold change in ARG abundance across drug classes for individual antibiotic enrichment treatments relative to FD. Fold change was calculated as the ratio of mean CPM in each enriched group to mean CPM in FD for each drug class, pooled across both aerobic and anaerobic oxygen conditions and all three antibiotic concentration levels (1X, 10X, and 100X size-adjusted MIC). Blue circles represent the mean fold change across all antibiotic enrichment treatments combined; grey circles represent fold change for each individual antibiotic treatment (Amp = Ampicillin; Cef = Ceftiofur; Ery = Erythromycin; Lin = Lincomycin; Tet = Tetracycline; Van = Vancomycin).

### CEMG enhances detection of ARGs associated with diverse predicted resistance mechanisms

The FD ARG profile was dominated by antibiotic inactivation (46.15%) and target protection (39.56%), driven primarily by lincosamide nucleotidyltransferases (LNU; 44.14%) and *msr*-type ABC-F ribosomal protection proteins (39.56%), respectively (Fig. S5A & S5B). Culture enrichment revealed a pronounced oxygen-dependent restructuring of these profiles. Under anaerobic conditions, both CEMG-NoAB and CEMG-AB showed enrichment of efflux- and target protection-dominated ARG profiles, with MFS efflux pumps as the leading gene family (31.67% and 40.37%, respectively) and efflux accounting for 44.06% and 45.84% of total resistance mechanisms (Fig. S5A & S5B). In contrast, aerobic enrichment favored antibiotic inactivation as the dominant mechanism (CEMG-NoAB: 48.79%; CEMG-AB: 38.99%), with *ant*(4’) aminoglycoside nucleotidyltransferases (10.58% and 12.43%) and *erm* 23S rRNA methyltransferases (11.24% and 7.50%), and *cfr* 23S rRNA methyltransferases (10.16% and 7.14%) - both *erm* and *cfr* conferring resistance through ribosomal target alteration - among the most abundant gene families alongside MFS efflux pumps (Fig. S5A & S5B).

### ARG-MGE linkage and co-selection of ARGs and heavy metal resistance gene

While reference-based analysis captured broad shifts in resistome composition following culture enrichment, it provided limited insight into microbial host associations and genomic context of ARGs, particularly linkages to MGEs. Hybrid assembly of short- and long-read data helped resolve these associations. Overall, hybrid assembly identified a total of 784 ARGs across 651 unique contigs, representing 100 unique ARGs. Of these, 59.3% (465/784) were co-localized with at least one MGE class across 353 ARG-MGE bearing contigs (Table S7). Plasmids were the most prevalent MGEs, identified on 230 contigs (65.2%), followed by insertion sequences (116 contigs, 32.9%), ICEs and IMEs (77 contigs, 21.8%), integrase/resolvase signatures (59 contigs, 16.7%), and phage or viral sequences (15 contigs, 4.2%) (Fig. 5A). Co-occurrence of multiple MGE classes on single contigs was frequently observed. Seventy contigs carried both insertion sequence (IS) elements and plasmid signatures, while 20 contigs exhibited triple co-occurrence of IS elements, integrase/resolvase genes, and plasmid classifications. These composite genetic architectures suggest mosaic MGE structures and indicate layered potential for horizontal transfer of ARGs within these genomic contexts (Fig. 5B). Plasmids accounted for the highest total ARG-MGE associations, spanning 65 unique ARGs across all major drug classes, while insertion sequences flanked 29 unique ARGs within 2 kb, with aminoglycoside, phenicol, lincosamide, and β-lactam resistance genes most frequently represented. ICEs and IMEs were associated with 20 unique ARGs, with tetracycline resistance genes (*tet*(M), *tet*(W), *tet*(Q), *tet*(36)) and MLS resistance genes (*erm*(F), *erm*(47)) showing the highest representation in this context.

**Figure 5.**
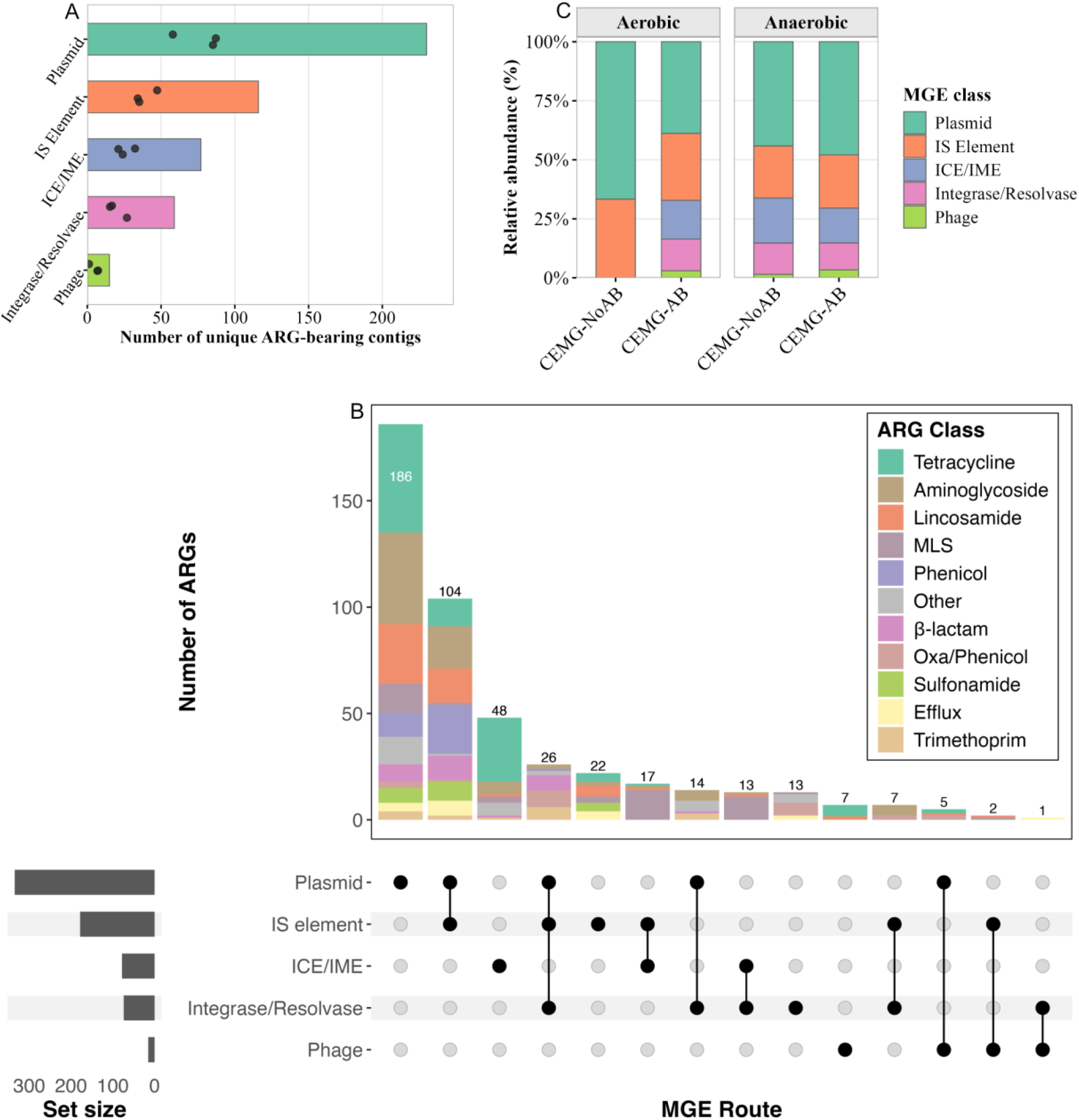
Diversity and composition of MGEs across assembled contigs and enrichment conditions. (**A**) Total number of unique ARG-bearing contigs associated with each MGE class. Bars represent the overall count summed across all replicates and sample groups; dots indicate per-replicate counts. MGE classes are ordered from most to least prevalent. (**B**) UpSet plot showing the number of ARG instances (y-axis, bars) distributed across combinations of MGE route classes (x-axis, filled dots and connecting lines), stratified by drug class (bar fill). Only intersections containing at least one ARG instance are shown. Set sizes (bottom left) indicate the total number of ARG instances associated with each MGE route class regardless of co- occurrence. **(C)** Relative abundance (%) of MGE classes among ARG-bearing contigs within each enrichment group, faceted by oxygen condition with CEMG-NoAB and CEMG-AB shown as separate bars within each facet. Each bar is normalized to 100% of MGE-associated contigs within that group.

Plasmid-associated ARG-bearing contigs were the most abundant across all enrichment conditions, accounting for 39-67% of ARG-MGE associations within each group (Fig. 5C). ICE/IME-associated contigs were absent from aerobic CEMG-NoAB but represented 14.8- 19.1% of associations under anaerobic CEMG-AB, suggesting that microbial hosts carrying conjugative transfer potential are selectively enriched under antibiotic enrichment and anaerobic conditions (Fig. 5C).

Co-localization analysis of ARG-MGE-bearing contigs further revealed the presence of biocide and metal resistance genes, with profiles varying markedly by oxygen condition (Fig. S6A). FD samples were dominated by sodium (44.4%), arsenic (33.3%), and zinc (22.2%) resistance, corresponding to the genes *pcm*, *arsM*, and *troB*, respectively (Fig. S6B). Under anaerobic enrichment, both CEMG-NoAB and CEMG-AB displayed diverse biocide and metal resistance profiles spanning zinc (21.4% and 17.1%), arsenic (13.4% and 12.3%), triton (12.6% and 9.2%), cadmium (7.2% and 9.5%), tetraphenylphosphonium, spermidine, sodium, and acriflavine, with a substantial “Other” fraction (21.2% and 23.3%) reflecting additional resistance classes beyond the top ten (Fig. S6A). In contrast, aerobic enrichment produced a markedly more restricted profile. Aerobic CEMG-NoAB was composed exclusively of cetrimide and triclosan resistance (each 50%), while aerobic CEMG-AB was similarly dominated by cetrimide (50.1%) and triclosan (34.7%), with an additional contribution from tetraphenylphosphonium resistance (15.3%). At the gene level, aerobic enrichment was characterized by *norA* and *smeT* in CEMG- NoAB, and *norA*, *smeT*, *qacF*, and *lmrS* in CEMG-AB, reflecting a shift toward biocide efflux-associated genes under aerobic conditions (Fig. S6B).

### ARG-MGE-host linkage

Taxonomic classification of ARG-bearing contigs using Kraken2 yielded classifications for 569 of the 651 ARG-bearing contigs identified in the hybrid assemblies. Among them, the majority were affiliated with *Bacillota* (304 contigs; 53.4%), followed by *Pseudomonadota* (168; 29.5%), *Bacteroidota* (48; 8.4%), and *Actinomycetota* (25; 4.4%), with the remaining 24 contigs (4.2%) assigned to other phyla (Fig. S7A). The overall abundance of ARG-MGE-associated contigs was higher in both CEMG-NoAB (*p* < 0.05) and CEMG-AB (*p* < 0.01) compared to FD (Fig. S7B).

The four-level linkage diagram linking drug class, ARG gene, bacterial genus, MGE class revealed distinct host associations and mobilization contexts across drug classes (Fig. 6, Table S8). Aminoglycoside resistance genes - including *aac(6’)-I*, *aac(6’)-lid*, *aadA1*, *aph(3”)-Ib*, and *aph*(*6*)*-Id* - represented the largest resistance drug class block and were distributed across *Acinetobacter*, *Enterococcus*, *Escherichia*, and *Hafnia*, with plasmid co-localization predominating. Tetracycline resistance genes displayed the broadest host associations, spanning *Streptococcus*, *Proteus*, *Enterococcus*, *Citrobacter*, and *Oscillibacter*, and were linked to a diverse array of MGE contexts including ICE/IME, IS elements, and plasmids. Key genes in this class included *tet*(M), *tet*(W), *tet*(H), *tet*(J), *tet*(L), and *tetB*(P). MLS resistance genes - primarily *erm*(F), *erm*(47), *erm*(D), and *mef*(En2) - were predominantly associated with *Streptococcus*, *Peptostreptococcus*, and *Vagococcus*, and were frequently co-localized with ICE/IME and IS elements, suggesting integrative mobility as a key dissemination route for this class. Lincosamide resistance gene (*lnu*(C)) was distributed across *Streptococcus*, *Jeotgalibaca*, and *Oscillibacter*, primarily in plasmid and IS element contexts. β-lactam resistance genes (*blaCMY*, *blaCMY-181*, *blaACC-1a*, *blaBCL*) were concentrated in *Escherichia*, *Citrobacter*, *Comamonas*, and *Acinetobacter*, and were almost exclusively plasmid-associated, often alongside IS elements. Oxazolidinone/phenicol resistance (*poxtA2*) was detected in *Enterococcus*, *Bacillus*, and *Streptococcus*, with co-localization spanning IS elements, plasmids, and ICE/IME contexts. Sulfonamide (*sul2*), trimethoprim, and efflux (*mdtM*) resistance genes were largely confined to *Escherichia* and *Citrobacter* and were predominantly plasmid-associated.

**Figure 6.**
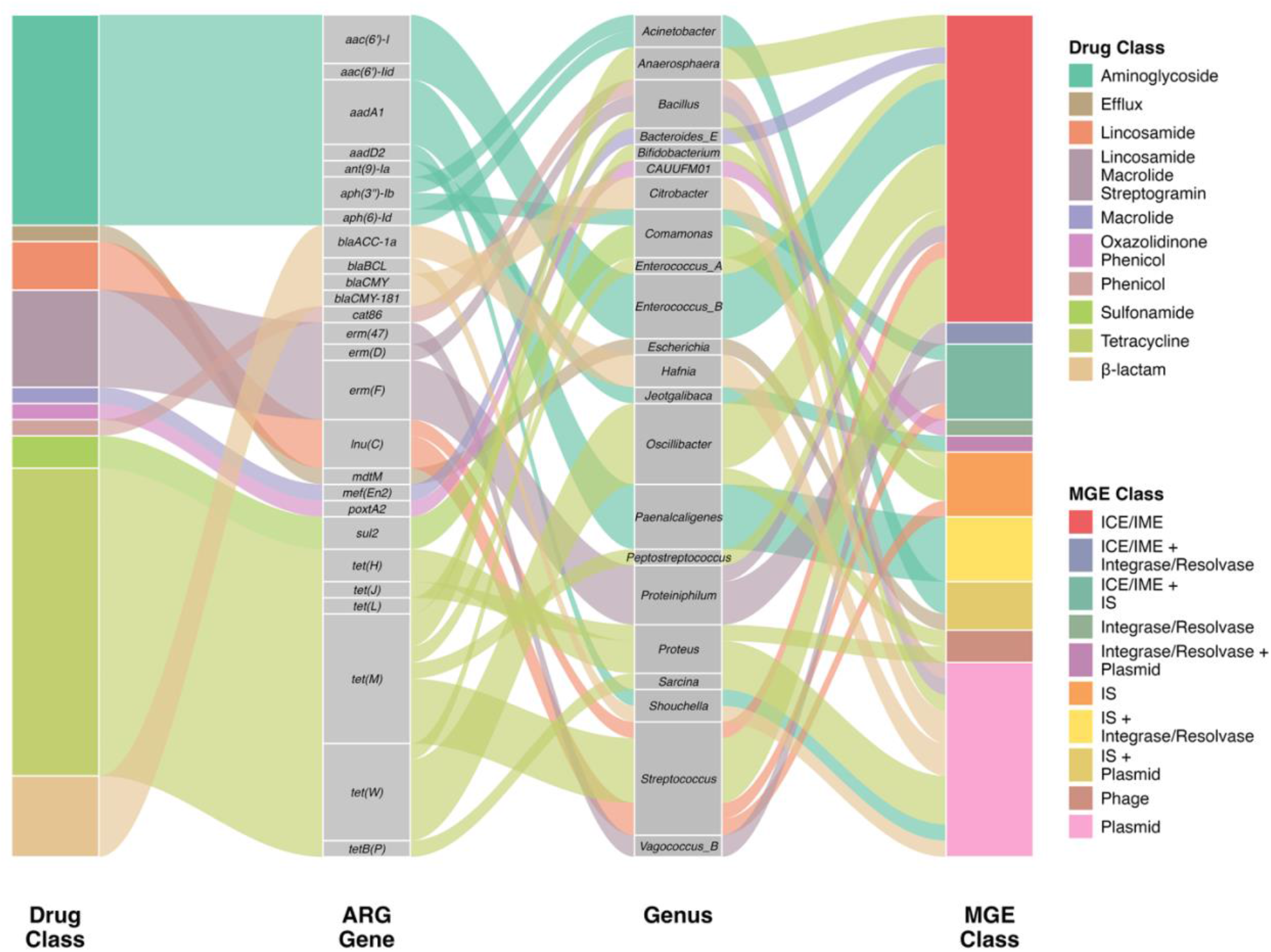
Co-occurrence patterns of ARGs, host genera, and MGEs. Sankey diagram showing co-occurrence patterns among resistance drug classes (left), ARG genes, bacterial genera, and MGE classes (right), based on contig-level co-localization. Flow width is proportional to the number of ARG-bearing contigs sharing each drug class-gene-genus-MGE combination.

## Discussion

Our results suggest that culture-enriched metagenomics not only improves the detectability of ARGs but also actively reshapes the observed resistome by selectively amplifying specific microbial populations, consistent with previous reports (9, 30). Unenriched digestate samples showed a comparatively homogeneous community structure, substantial representation of poorly resolved taxa affiliated with phyla characteristic of stable anaerobic digestion communities (28, 29). By contrast, enrichment reduced the dominance of this baseline community and favored bacterial groups better adapted to laboratory cultivation. This pattern reinforces an important conceptual distinction: direct metagenomics is better suited to describing the *in-situ* community within the digester (31), whereas culture enrichment preferentially captures the cultivable fraction that may be more relevant to post-digestion survival, dissemination, and hazard assessment.

The enrichment-driven expansion of species-level diversity, particularly the recovery of many species uniquely detected in antibiotic-supplemented cultures, is especially noteworthy from a surveillance perspective. Previous studies have also reported increased detection of clinically relevant ARGs and taxa following selective enrichment (19). The appearance of taxa such as *E. dispar, E. casseliflavus, E. hormaechei, C. amalonaticus,* and *A. insolitus* following antibiotic enrichment, organisms with recognized clinical and environmental relevance (32–35), suggests that these populations persist in digestate at low abundance or in a suppressed state, becoming detectable only under selective antibiotic cultivation. From a One Health perspective, this supports the idea that direct shotgun sequencing alone can underestimate the diversity of viable, resistant, and potentially transmissible bacteria present in environmental samples like digestate.

This enrichment-driven shift was also reflected in a 10-fold increase in overall ARG abundance, consistent with observations from wastewater systems (19). Importantly, this shift was not limited to a greater ARG burden but also involved a broadening of the ARG drug-class profile. Fresh digestate was dominated by ARGs associated with MLS resistance, whereas enriched communities displayed a broader profile across tetracycline, β-lactam, aminoglycoside, phenicol, rifamycin, glycopeptide, and other resistance classes. Although a broad range of ARG classes have been reported in manure and digestate (31, 36), the strong dominance of MLS resistance in our unenriched digestate likely obscured less abundant resistance determinants that became more detectable following culture enrichment. Notably, several clinically important ARG families were recovered exclusively in enriched samples, including genes associated with glycopeptide resistance, β-lactamase genes of the CTX-M, OXA, and TEM families, some variants of which are known to confer extended-spectrum β-lactamase activity, though variant-level confirmation was not performed in the current study. The *cfr* 23S rRNA methyltransferase is particularly notable given its cross-class activity spanning phenicols, lincosamides, oxazolidinones, pleuromutilins, and streptogramin, meaning a single gene simultaneously undermines multiple antibiotic classes used in both human and veterinary medicine. These findings align with a growing body of evidence that the cultivable, low-abundance fraction of environmental resistomes carries disproportionate clinical relevance relative to its abundance (37–39).

The strong representation of tetracycline resistance following multiple enrichment treatments is also of importance. Tetracycline resistance genes are widely distributed in livestock-associated bacteria and are often associated with MGEs, which may facilitate persistence even in the absence of tetracycline (39–41). The fact that tetracycline resistance increased across several antibiotic enrichments, rather than just with tetracycline, likely reflects co-selection and co-maintenance of multidrug resistance determinants in the same bacterial population or linked mobile elements (42). A similar interpretation may explain the increases in cephalosporin, aminoglycoside, phenicol, glycopeptide, and diaminopyrimidine resistance observed under non-cognate antibiotic treatments.

The generally weak effect of increasing antibiotic concentration on community diversity and ARG richness suggests that, once resistant populations are selected, further concentration increases do not broaden recovery and may instead narrow the cultured community. This supports the use of the size-adjusted lowest MIC for broad AMR surveillance, as higher concentrations may reduce the ecological representativeness of enriched communities.

The predominance of plasmid- and ICE/IME-associated ARGs across all major drug classes has direct implications for post-digestion dissemination risk. MGEs are the primary vehicle for cross-species ARG transfer, so, even if anaerobic digestion reduces the abundance of resistant organisms, the persistence of ARGs on conjugative and replicative elements sustains horizontal transfer potential upon land application of digestate - independent of the organisms that originally carried them (43). The co-localization of ARGs with biocide and metal resistance genes on the same contigs further supports co-selection as a plausible mechanism maintaining AMR in post-digestion environments. The high relative abundance of zinc and arsenic resistance genes is consistent with the widespread use of zinc-based compounds as feed additives in food animal production (44), and with previous reports of zinc co-selecting for antibiotic resistance through shared genetic linkage on MGEs (45). Because antibiotic and heavy metal resistance determinants are frequently co-located on the same mobile element, metal selection pressure in manure-amended soils may indirectly sustain antibiotic resistance gene pools long after antimicrobial use ceases.

A key strength of this study is the ability to link ARGs and their MGE contexts to putative bacterial hosts at the contig level, a resolution that read-based approaches cannot provide. Particularly significant is the broad host distribution of tetracycline resistance ARGs across phylogenetically diverse genera - spanning both Gram-positive commensals such as *Streptococcus* and *Vagococcus* and Gram-negative opportunists such as *Proteus*, *Citrobacter*, and *Escherichia* - which, combined with frequent ICE/IME co-localization, suggests that conjugative dissemination of tetracycline resistance is not confined to a single host lineage but is distributed across the enriched community. This observation reinforces the broader point that culture enrichment can surface clinically relevant ARG-MGE-host combinations that remain invisible in bulk metagenomic profiles, and that the cultivable fraction recovered under selective conditions may disproportionately represent the transmissible, high-risk segment of the environmental resistome.

The distribution of ARG-MGE-host combinations also differed markedly between aerobic and anaerobic enrichments, indicating that oxygen availability is a primary determinant of which resistance determinants are recovered. Previous studies have also shown that oxygen exposure can restructure microbial communities and alter ARG profiles (46). This suggests that, in digestate-derived enrichment systems, redox conditions may act as a first-order ecological determinant that governs which bacterial groups proliferate and, consequently, which resistance mechanisms are detectable. Aerobic enrichment favored communities dominated by *Alcaligenes* and related taxa, whereas anaerobic enrichment selected for *Proteus*, *Streptococcus*, *Vagococcus*, and *Fermentimonas*. These shifts likely reflect major physiological differences in respiratory capacity, substrate use, and competitive fitness under oxic versus anoxic conditions (47), with corresponding changes in the resistome driven by shifts in host community structure. Incorporating both aerobic and anaerobic enrichment conditions may therefore improve AMR surveillance, as some ARGs and their associated hosts may remain at low abundance under anaerobic conditions but become more detectable under aerobic conditions, enabling identification of clinically relevant taxa that may proliferate following land application of manure or digestate.

While these findings provide important insights into the structure and mobilization potential of the resistome, several limitations should be considered when interpreting the results. Firstly, not all microorganisms present in digestate can grow under the culture conditions used in this study (48–50). Consequently, taxa that are unculturable under these conditions would remain undetected regardless of their abundance in the source material. Secondly, the use of a single culture medium likely constrained the diversity of organisms recovered, whereas employing a broader range of selective and non-selective media could capture a wider spectrum of microbial diversity (49). Thirdly, the observation that a substantial fraction of ARG instances lacked detectable MGE associations raises an important interpretive caution: the absence of detectable linkage in assembly-based analyses does not confirm chromosomal localization, as contig fragmentation and database gaps can obscure genuine mobile contexts. This is particularly relevant for glycopeptide resistance ARGs, which were absent from MGE-associated contigs despite being detected at the read level, leaving their genomic context unresolved (51). Finally, although the present workflow identifies ARG-MGE-host linkages and co-occurrence patterns consistent with mobilization potential, it does not directly quantify HGT frequency or confirm the expression of resistance phenotypes.

### Conclusions

Overall, this study demonstrates that culture-enriched metagenomics can substantially enhance AMR surveillance resolution in livestock digestate. By selectively amplifying viable microbial populations, CEMG enabled the recovery of low-abundance ARG-carrying taxa, expanded the detectable diversity of ARGs, and facilitated the identification of ARG-MGE-host co-localization patterns and ARG-metal co-selection signals that were largely obscured in direct metagenomic profiles. These findings indicate that assessments based solely on bulk community metagenomes may underestimate the diversity and mobilization potential of ARGs present in digestate. Integrating enrichment-based approaches with direct metagenomic analyses may therefore provide a more comprehensive framework for evaluating the environmental dissemination risk of AMR associated with livestock-derived anaerobic digestate.

## Methods

### Sample collection

Digestate was obtained directly from a continuously stirred (120 rpm), mesophilic (35°C) anaerobic digester (CSTR; New Brunswick Scientific BioFlo 110). The digester was operated in our laboratory for more than six years as a 14 L fed-batch system, with a working volume of 8 L and a solids retention time of 30 days. During the study period, the digester was in a stable operational state, with an average methane yield of 119.816 mL g^-1^ volatile solids (VS) Day^-1^ and a pH of 7.43. The reactor was fed with manure collected from a family-owned commercial dairy farm (Rosser Holsteins Dairy Farm, Rosser, Manitoba) with approximately 600 cows. It was fed approximately three times per week (e.g., Monday, Wednesday, and Friday), with 0.622 L of digestate removed and replaced with an equal volume of fresh manure, maintaining an 8 L working volume and a 30-day hydraulic retention time.

The collected digestate was divided into three aliquots for: (i) anaerobic culture enrichment within an anaerobic chamber, (ii) aerobic culture enrichment, and (iii) direct shotgun metagenomic sequencing (FD). Aliquots designated for anaerobic culture were immediately transferred in 50 ml Falcon tubes into an anaerobic chamber. Sampling was repeated monthly from mid-November to mid-January to obtain three biological replicates from the same digester.

### Anaerobic chamber conditions

All anaerobic procedures were performed in an anaerobic chamber (Bactron Anaerobic Chamber, Model: BACTRON600), maintained at 37°C using a built-in incubator. The chamber atmosphere was composed of 5% CO_2_, 5% H_2_, and 90% N_2_, supporting obligate anaerobe conditions. Anaerobiosis was regularly verified using anaerobic indicators (Oxoid Ltd., Hants, UK). All media and equipment were transferred into the chamber one day prior to sample collection.

### Culture media preparation and antimicrobial supplementation

Brain Heart Infusion (BHI) agar (Difco^TM^, Sparks, MD, USA) was used as the general enrichment medium in this study, supplemented with 1 mg/L Vitamin K, 10 mg/L Hemin and 0.5 g/L L-cysteine to promote the growth of obligate anaerobes. To enrich antimicrobial-resistant communities, six antimicrobials representing major antibiotic classes were incorporated individually into separate BHI agar plates, with each plate containing only one antimicrobial at a given concentration. Five classes were selected on the basis of their documented use in livestock production and prior detection of associated ARGs in dairy manure and anaerobic digestate processed in the same laboratory digester system (31): ampicillin (β-lactam; 0.4 µg/100 ml), tetracycline (tetracycline; 1.6 µg/100 ml), lincomycin (lincosamide; 1.6 µg/100 ml), erythromycin (macrolide; 0.8 µg/100 ml), and ceftiofur (third-generation cephalosporin; 0.05 µg/100 ml). Vancomycin (glycopeptide; 12.5 µg/100 ml) was additionally included given its critical importance as a last-resort antibiotic in human medicine and prior evidence of glycopeptide resistance genes in digestate from the same digester (31). Antibiotic concentrations were determined using size-adjusted lowest minimum inhibitory concentrations (MICs) following Bengtsson-Palme *et al.* (2016) (52). These values represent the estimated upper limit of the minimal selective concentration (MSC), defined as the lowest antibiotic concentration at which resistant bacteria gain a selective advantage over susceptible populations in environmental samples (52).

### Inoculation and incubation

Within the anaerobic chamber, digestate samples were diluted to 10-fold (10^-1^) in sterile BHI broth (Difco^TM^, Sparks, MD, USA) supplemented with 0.05% L-cysteine. A 200 µL aliquot of the diluted sample was spread onto each prepared agar plate using sterile glass beads. Anaerobic plates were incubated for 5 days at 37°C within the chamber. Simultaneously, an equivalent set of plates was inoculated aerobically in a biosafety cabinet (Thermo Electron Corp.) and incubated for four days at 37°C. In total, seven treatment groups were prepared under both anaerobic and aerobic conditions, comprising one non-antibiotic enrichment group (CEMG- NoAB) and six antibiotic-supplemented enrichment groups (collectively referred to as CEMG- AB). Within CEMG-AB, each antimicrobial was applied at three concentrations (1X size-adjusted lowest MIC, 10X size-adjusted lowest MIC, and 100X size-adjusted lowest MIC) to assess dose-dependent effects on anaerobic communities; this dose-response design was restricted to anaerobic enrichment. Following incubation, visible colonies were scraped and pooled by treatment group using 2 mL sterile BHI broth per plate. Colonies from the anaerobic plates with different antibiotic concentrations were pooled separately to retain dose-dependent resolution. All pooled samples were subsequently used for DNA extraction and downstream molecular analyses.

### DNA extraction and sequencing

DNA was extracted from fresh digestate (FD) and CEMG samples using an EZNA Soil DNA kit (Omega Bio-Tek, Inc.) as per the manufacturer’s protocol. A total of 250 µL of fresh digestate or scraped colony biomass resuspended in BHI broth was used as input for each extraction. Illumina sequencing libraries were prepared using the SparQ DNA Frag & Library Prep kit with UMI (Unique Molecular Identifier) indexing (Quantabio, Beverly, MA, USA). Libraries were pooled at equimolar concentrations and size-selected using SparQ PureMag beads (dual-size selection). Sequencing was performed on an Illumina NovaSeq 6000 generating (2 × 150 bp) paired end reads.

For long-read sequencing, DNA was cleaned using the ProNex® Size-Selective Purification System (NG2001; Promega, Madison, WI, USA) with protocol 6A at a 1x (v/v) bead ratio to remove contaminants and low-molecular-weight dsDNA, with a minimum input of 200 ng gDNA per sample (53). Nanopore libraries were prepared using the Rapid Barcoding Kit 96 V14 (SQK-RBK114.96; Oxford Nanopore Technologies, Oxford, UK) and loaded onto a PromethION R10.4.1 flow cell (FLO-PRO114M) following the manufacturer’s flow cell priming and loading protocol. Base-calling was performed using Dorado (v1.3.2, HAC v4.3.0 model).

### Initial processing of the Illumina and Nanopore reads

Raw Illumina paired-end reads were adapter-trimmed and quality-filtered using fastp v1.0.1 (54) with TruSeq adapter sequences specified explicitly (Read 1: AGATCGGAAGAGCACACGTCTGAACTCCAGTCA; Read 2: AGATCGGAAGAGCGTCGTGTAGGGAAAGAGTGT), polyG tail trimming enabled, and default quality filtering parameters applied. Sequencing quality metrics were assessed using MultiQC v1.30 (55).

Host and plant-derived sequence removal was performed separately for Illumina and Nanopore reads using Minimap2 (v2.30-r1287) (56) against the same six reference genomes: *Zea mays* (NCBI accession no. GCF_902167145.1), *Triticum aestivum* (NCBI accession GCF_018294505.1), *Medicago sativa* (NCBI accession GCA_051527215.1), *Hordeum vulgare* (NCBI accession GCF_904849725.1), *Brassica napus* (NCBI accession GCF_020379485.1), and *Bos taurus* (NCBI accession GCF_002263795.3). For Illumina reads, host removal was applied sequentially, mapping paired-end reads against each reference genome in turn using the short- read preset (-x sr, --secondary=no); reads with a mapping quality score ≥ 30 to primary alignments were identified as host-derived and removed using filterbyname.sh (BBMap suite), with unmapped reads from each step carried forward as input to the next. For Nanopore reads, all six reference genomes were concatenated into a single combined reference, indexed once with Minimap2, and reads were aligned using the ONT-specific preset (-x map-ont); only reads that failed to map to the combined reference (SAM flag -f 4) were retained. No additional quality trimming was applied to Nanopore reads prior to host removal, as base-calling and adapter trimming were performed upstream by the PromethION basecaller during signal processing. Host-decontaminated reads from both platforms were used for all subsequent analyses.

### Read-based analysis

Quality-controlled paired-end Illumina reads were profiled for ARGs using the Resistance Gene Identifier (RGI v6.0.5) with the BWT workflow, aligning reads against the Comprehensive Antibiotic Resistance Database (CARD v4.1.1) using KMA (v1.6.6) as the aligner, including wildcard variants (57). Only hits with ≥90% read coverage and ≥98% nucleotide identity of the reference sequence were retained. ARG abundance was quantified as counts per million total sequencing reads (CPM), calculated as:

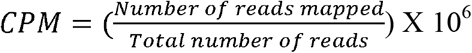

For taxonomic profiling, reads were classified at the species level using MetaPhlAn v4.2.2 (58) with the CHOCOPhlAnSGB vJan25 marker gene database, with read alignment performed internally using Bowtie2 (v2.5.4) (59) to determine shifts in microbial communities between fresh digestate and enriched samples.

### Hybrid metagenome assembly

Hybrid metagenome assemblies were generated by combining quality-controlled Illumina paired-end reads with Nanopore long reads using SPAdes (v4.2.0) in metagenomic mode (60). Illumina reads were grouped by antimicrobial type, with all three concentration levels co-assembled within each antimicrobial-defined group to maximize contig length and ARG-MGE resolution. Nanopore long reads were incorporated during assembly using the --nanopore option. Default k-mer settings were applied, and assemblies were performed using the SPAdes internal error-correction and iterative assembly pipeline. Contig-level taxonomic classification of hybrid-assembled contigs was performed using Kraken2 (v2.17.1) (61) against the PlusPFP database (20250402); only contigs ≥1 kb that received a taxonomic classification were retained for downstream ARG-MGE-host linkage analyses. MetaPhlAn was used for read-based community profiling given its clade-specific marker gene approach and high species-level accuracy in complex environmental samples, while Kraken2 was applied separately for contig-level taxonomic classification to support ARG-MGE-host linkage analyses.

### Functional annotation and construction of ARG-MGE co-localization

ARGs were identified from assembled contigs using two complementary tools. RGI (v6.0.5) (57) was run in contig mode (--input_type contig) against a local CARD database (v4.1.1) using BLAST as the alignment algorithm, with loose, Strict, and Perfect hit categories all reported (-- include_loose). AMRFinderPlus (v4.2.5) (62) was run in nucleotide mode (-n) against the NCBI Reference Gene Catalog with the extended --plus database enabled, applying minimum nucleotide identity and coverage thresholds of 90% and 70%, respectively. ARG calls detected by both tools on the same contig were retained for all downstream analyses.

Protein-coding genes were predicted on ARG-bearing contigs using Prodigal (v2.6.3) in metagenomic mode (-p meta) (63) to provide a consistent coordinate space for MGE annotation. MGEs were then annotated using a multi-tool pipeline. Plasmids were identified using geNomad (v1.12.0) (64) in end-to-end mode against the geNomad database, and by MOB-suite mob_recon (v3.1.9) (65) against the mob_suite reference database. A contig was classified as plasmid-associated if identified by either tool, with geNomad and MOB-suite serving as complementary callers to maximise detection sensitivity. Where a contig was classified as both a plasmid and a phage/virus, the plasmid classification took precedence over conjugative element classification to avoid double-counting. Phage and viral sequences were detected using geNomad and VirSorter2 (v2.2.4; executed in a pinned environment: Python 3.8, scikit-learn 0.22.1) (66) and quality-assessed with CheckV (v1.1.1) against checkv-db-v1.5 (67). Insertion sequences were identified using ISEScan (v1.7.3) (68) with default parameters. Site-specific recombinases (integrases and resolvases) were detected by scanning Prodigal (v2.6.3) (63)-predicted proteins against Pfam profiles PF00589 (phage integrase), PF00239 (resolvase), PF07508 (recombinase), and PF03432 (relaxase) using HMMER hmmsearch (v3.3.2) (69) at e-value < 1 × 10^-5^. Conjugative element classification was performed using a 129-profile HMM library combining the four above-mentioned Pfam profiles with 125 CONJScan T4SS profiles (CONJScan models v2.1.0; relaxase MOB families, type IV coupling protein, and mating-pair-formation components) (70), with signature-gene hits scored against ARG-flanking windows of ±2 kb for insertion sequences and recombinases, and ±25 kb for conjugative elements (ICEs and IMEs), following the approach described in Rahman *et al*. (2025) (71). Plasmid and phage classifications were assigned at the contig level. In cases where multiple MGE types were detected on the same contig, plasmid and phage calls took precedence over conjugative element classification. Biocide and metal resistance genes were identified on ARG-bearing contigs using Abricate (v1.0.4) (72) with the BacMet2 experimentally confirmed resistance gene database (built 2026-Apr-3; 746 protein sequences), applying minimum nucleotide identity and coverage thresholds of 90% and 90%, respectively.

To quantify the short-read abundance of ARG-associated genomic regions across enrichment conditions, assembled contigs were classified into co-localization categories - defined here as the presence of ARGs, MGEs, and/or biocide and metal resistance genes on the same contig based on genomic coordinates - using a custom Python script: ARG-only, MGE-only, BacMet-only, ARG-MGE co-localized, ARG-BacMet co-localized, MGE-BacMet co-localized, and ARG-MGE-BacMet co-localized contigs. Each category was extracted as a separate FASTA reference set. Quality-controlled host-filtered Illumina reads were mapped to these mapping reference sets using BWA-MEM (v0.7.18) (73) at a minimum nucleotide identity of 95%. Alignments were sorted and indexed using Samtools (v1.21) (74), and per-contig coverage was assessed using ‘samtools coverage’. Contigs were retained if they met minimum thresholds of ≥80% breadth of coverage, ≥80% query coverage, and ≥2x mean depth. Read counts for contigs passing these filters were summed and normalized to CPM (as defined in the Read-based analysis section) to enable cross-sample comparisons across enrichment conditions.

### Statistical analysis

All statistical analyses were performed in RStudio (version: 2025.09.0; R 4.5.2) using the vegan (2.7-1) (75), rstatix (0.7.2) (76), lme4 (1.1.37) (77), and emmeans (1.11.1) (78) packages. Bray-Curtis dissimilarity matrices were computed from genus-level relative abundance profiles and ARG compositional profiles using the vegdist function and visualized by principal coordinates analysis (PCoA). Differences in microbial community composition and resistome structure across treatment groups were tested using permutational multivariate analysis of variance (PERMANOVA; adonis2, 999 permutations), with effect sizes reported as R^2^ and significance determined at *p* < 0.05. Pairwise PERMANOVA between antibiotic treatment groups was performed using the same settings with Benjamini-Hochberg *p*-value adjustment. Homogeneity of multivariate dispersion was assessed using betadisper followed by permutation testing (permutest, 999 permutations) to confirm that observed compositional differences reflected true centroid separation rather than unequal within-group variability.

Alpha diversity was quantified using the Shannon index (vegan::diversity) and compared across enrichment groups, antibiotic treatments, and dose levels using the Kruskal-Wallis test, with pairwise post-hoc comparisons performed using the Dunn test with Benjamini-Hochberg false discovery rate correction (rstatix::dunn_test). ARG richness differences between FD and enriched groups were assessed using the Wilcoxon rank-sum test. Differences in ARG-MGE co- localization frequencies across enrichment conditions were assessed using a linear mixed-effects model, with pairwise group comparisons performed using estimated marginal means (emmeans package) and *p*-values adjusted using the Benjamini-Hochberg procedure. Dose-response relationships between antibiotic concentration and community diversity or ARG richness were evaluated by fitting linear and quadratic regression models, with model selection based on the Akaike Information Criterion (AIC); the linear model was preferred where it yielded a lower or comparable AIC relative to the quadratic alternative. ARG abundance was quantified as counts per million total sequencing reads (CPM), calculated as mentioned in the previous section. Fold changes in ARG abundance between groups were calculated as ratios of group mean CPM values. Resistance mechanism and drug class profiles were summarized as descriptive compositional proportions (percentage of total ARG-mapping reads within each group) without inferential testing. Statistical significance was determined at *p* < 0.05 for all tests.

## Declarations

## Ethics approval and consent to participate

Not applicable.

## Data availability

Raw metagenome sequences described in this study have been deposited in the NCBI Sequence Read Archive under BioProject accession: PRJNA1454115. Large supplementary data tables are deposited to Figshare:

Table S1: https://doi.org/10.6084/m9.figshare.32942219,

Table S3: https://doi.org/10.6084/m9.figshare.32942243

Table S4: https://doi.org/10.6084/m9.figshare.32942258

Table S5: https://doi.org/10.6084/m9.figshare.32942270

Table S7: https://doi.org/10.6084/m9.figshare.32942300.

Supplementary Tables S2, and S6 are provided in the supplementary file accompanying this manuscript.

## Code availability

All custom scripts, pipelines and workflows used in the analyses are included in the GitHub repositories: MAGnet (https://github.com/zisanurrahman/MAGnet) and CEMG_analysis_pipeline (https://github.com/zisanurrahman/CEMG_analysis_pipeline). Analyses were performed using RStudio (version: 2025.09.0; R 4.5.2) or Python (version 3.12).

## Author contributions

HD and NC conceptualized the study. HD, NR and ASMZR designed the computational framework and performed bioinformatics analyses. NR and ASMZR generated the figures and visualizations and drafted the manuscript. NR conducted the molecular microbiology work, including culture enrichment, DNA extraction and library preparation. HD supervised the project, provided guidance on both experimental and computational framework design, data interpretation, manuscript revision, and secured funding. All authors reviewed, edited, and approved the final manuscript.

## Acknowledgements

The authors thank Rosser Holsteins Dairy Farm, Rosser, Manitoba for providing manure for this study, and the Department of Biosystems Engineering, University of Manitoba, for providing access to the anaerobic digester facility.

