## Supplementary material for "Optimizing a Culture-Enriched Hybrid Metagenomics Pipeline to Assess the AMR Footprint of Livestock Manure in Anaerobic Digestate"

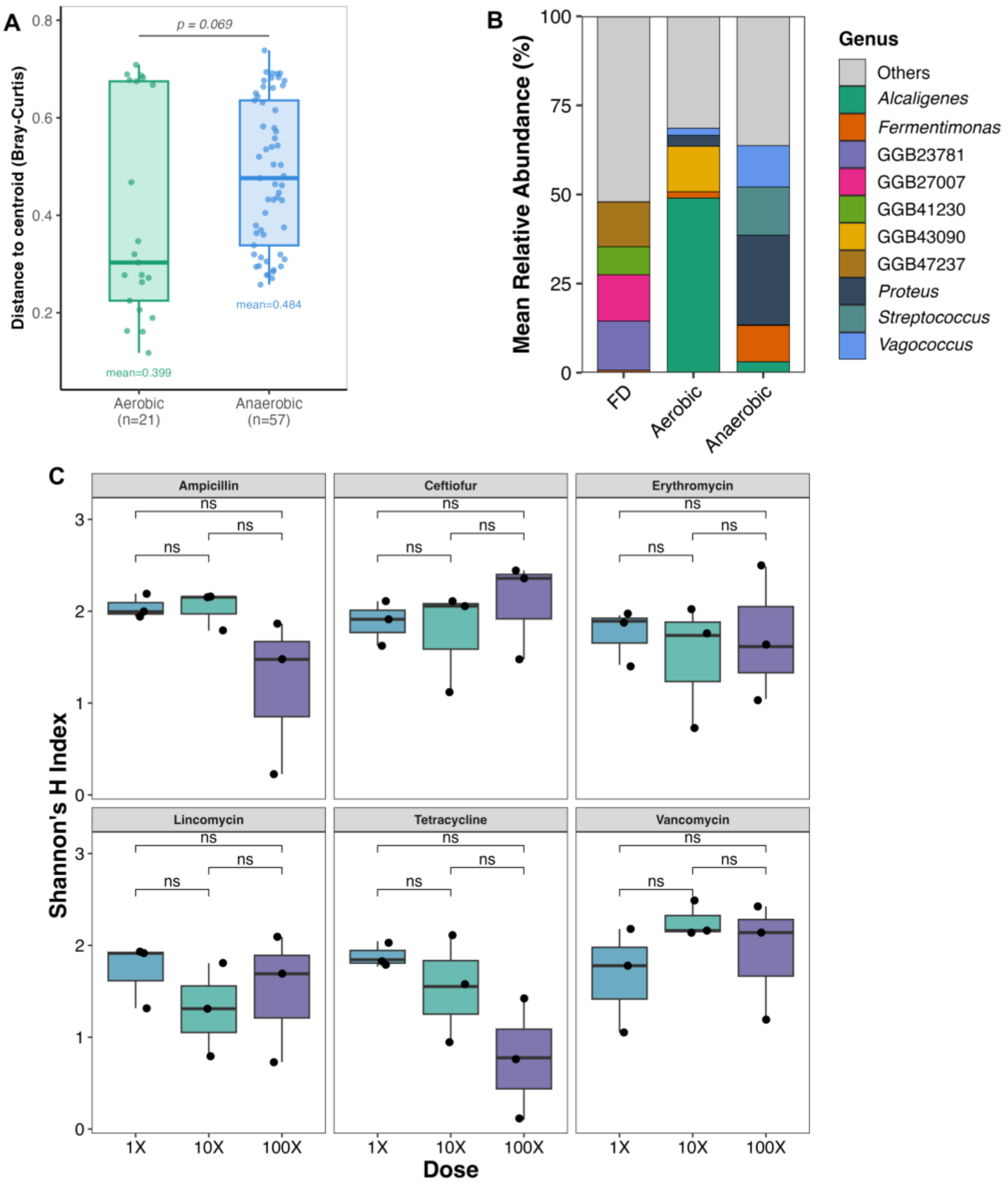


**Fig S1 Multivariate dispersion, community composition, and dose-dependent diversity patterns in digestate communities. (A)**Bray-Curtis distances to group centroid for aerobic and anaerobic culture-enriched communities, derived from the betadisper analysis. Individual data points represent samples; horizontal lines indicate group medians; boxes span the interquartile range. **(B)** Mean relative abundance of the top genera across FD, aerobic (pooled CEMG-NoAB and CEMG-AB), and anaerobic-enriched communities (pooled CEMG-NoAB and CEMG-AB). Genera outside the top displayed taxa are grouped as "Others"; GGB entries denote unclassified genome-based genus-level bins. **(C)** Shannon diversity index across three antibiotic concentration levels (1X, 10X, 100X the size-adjusted minimum inhibitory concentration) for each of the six antibiotic treatments, under anaerobic enrichment conditions. Data points represent individual biological replicates (n = 3 per dose level per antibiotic). Pairwise differences between dose levels were assessed using Dunn post-hoc tests (Benjamini-Hochberg correction) following Kruskal-

Wallis tests. Note that these pairwise comparisons differ from the linear dose-response models reported in Supplementary Table 4.

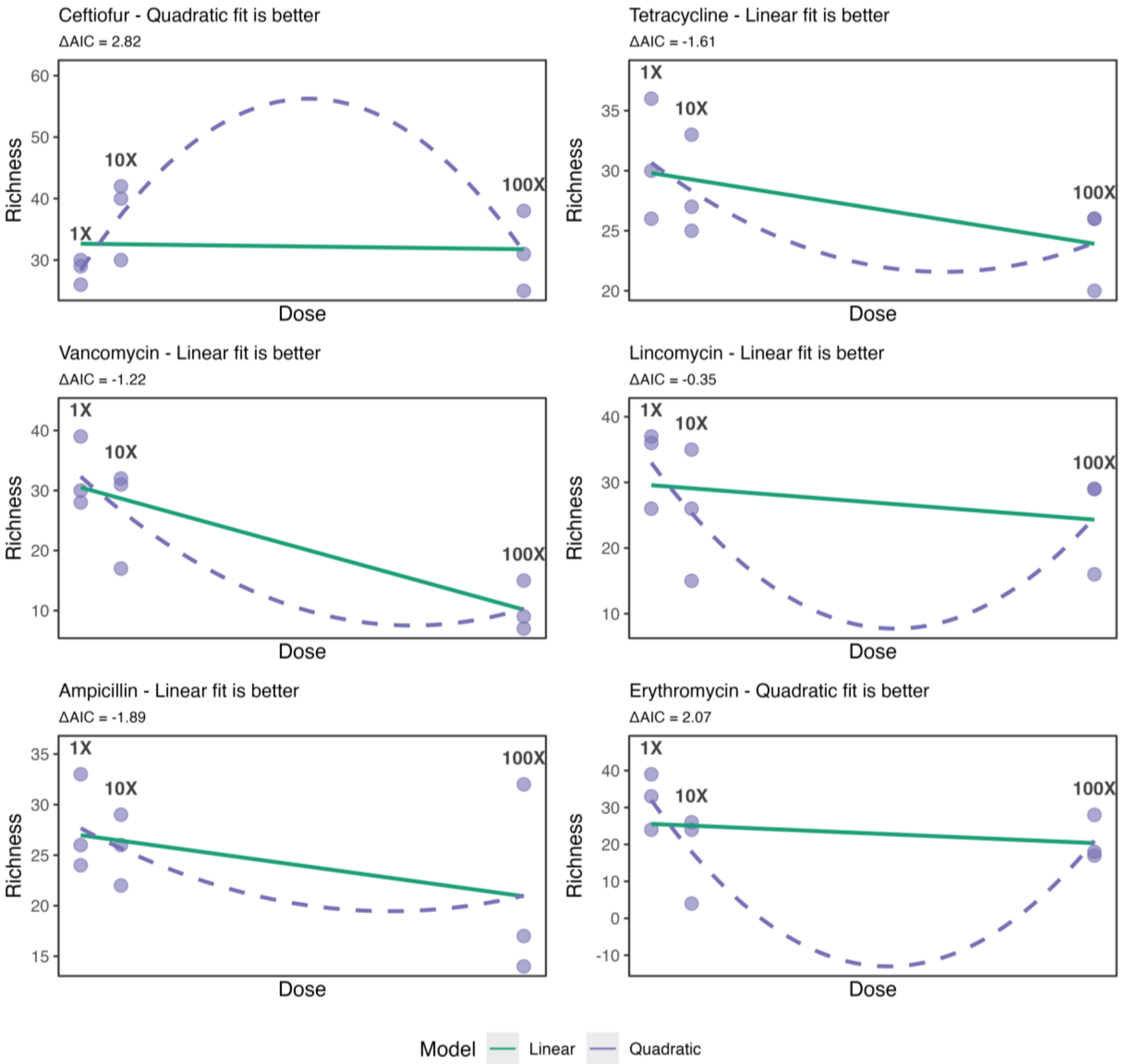


**Fig S2 Dose-response relationships of taxonomic richness under anaerobic antibiotic enrichment.** Taxonomic richness is shown across three dose levels (1X, 10X, and 100X) for each antibiotic treatment. Data points represent independent biological replicates (n=3 per dose). Solid lines indicate linear model fits, and dashed lines indicate quadratic model fits. Model selection is based on ΔAIC values shown within each panel.

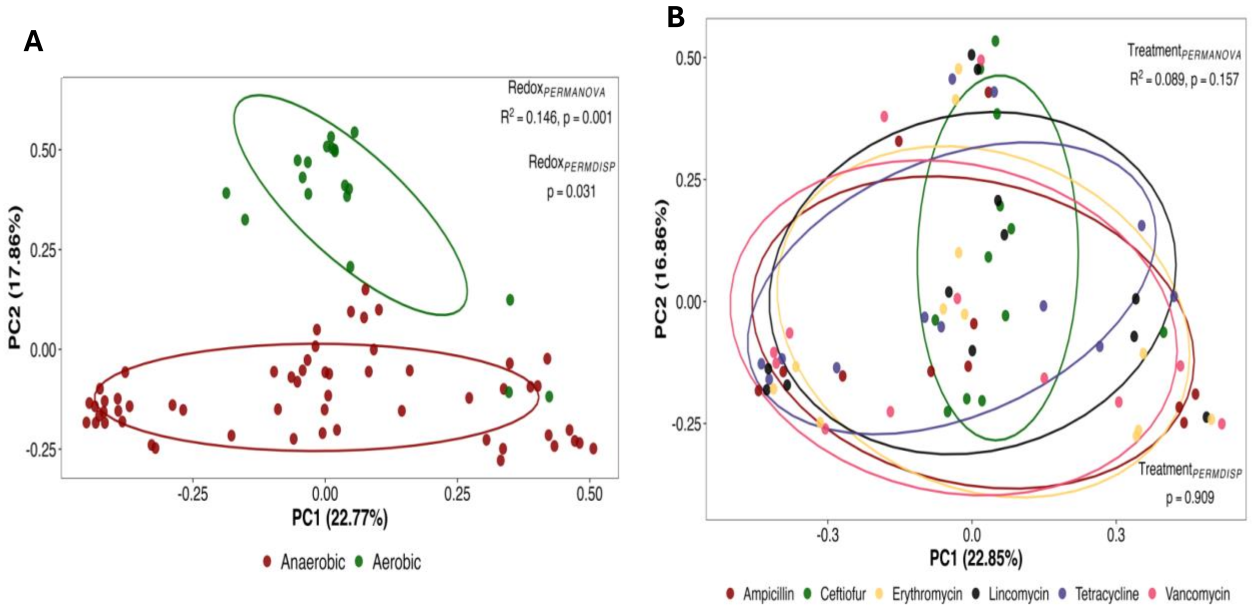


**Fig S3 Ordination of ARG composition.** (**A**) Principal coordinates analysis (PCoA) of ARG profiles based on Bray-Curtis dissimilarity, comparing aerobic and anaerobic enrichment conditions. Data points represent individual samples, and ellipses indicate group clustering. (**B**) PCoA of ARG profiles based on Bray-Curtis dissimilarity across antibiotic treatments. Data points represent individual samples, and ellipses indicate group clustering.

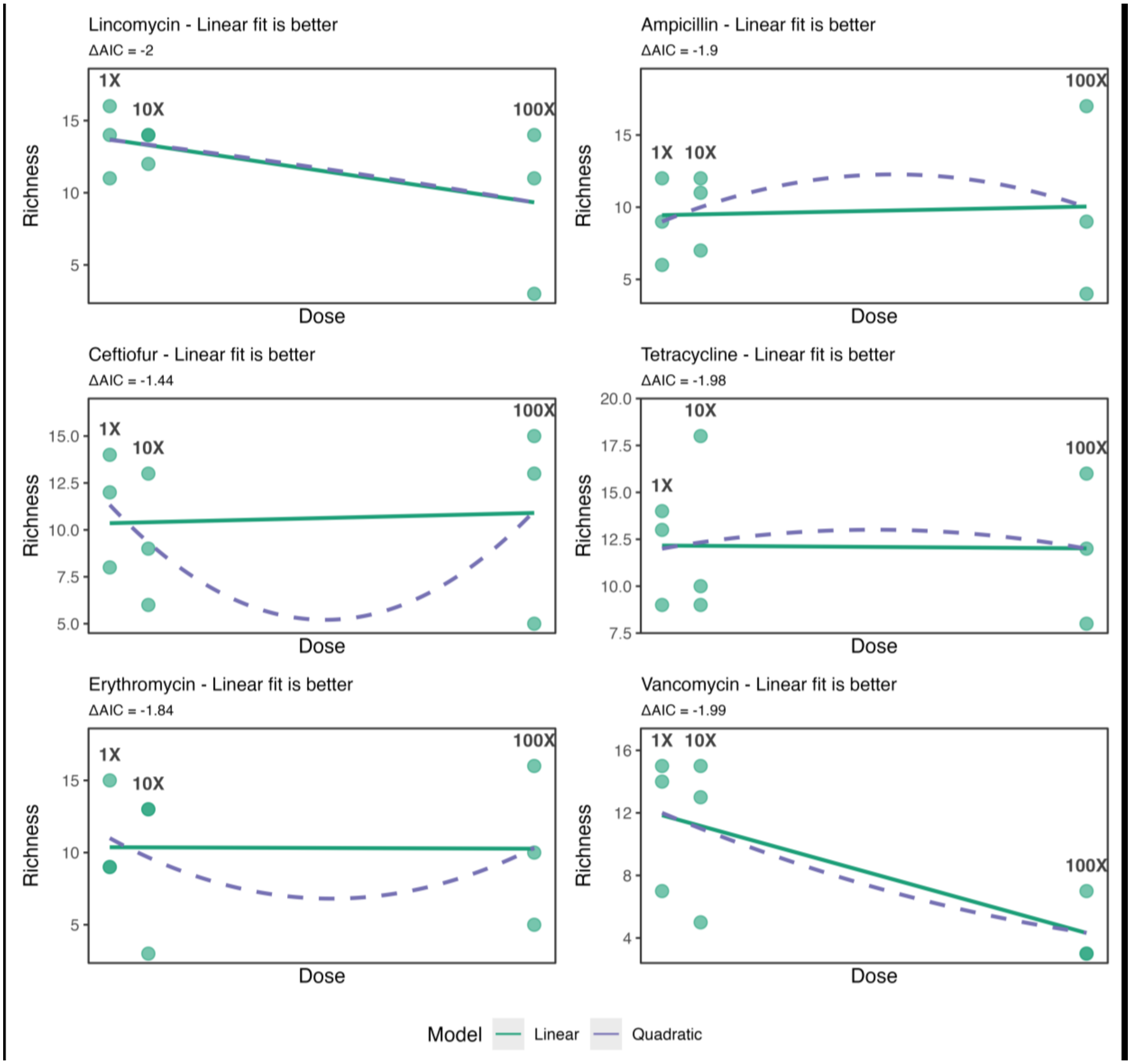


**Fig S4 Dose-response relationships of ARG richness under anaerobic antibiotic enrichment.** ARG richness is shown across three dose levels (1X, 10X, and 100X) for each antibiotic treatment. Data points represent independent biological replicates (n=3 per dose). Solid lines indicate linear model fits, and dashed lines indicate quadratic model fits. Model selection is based on ΔAIC values shown within each panel.

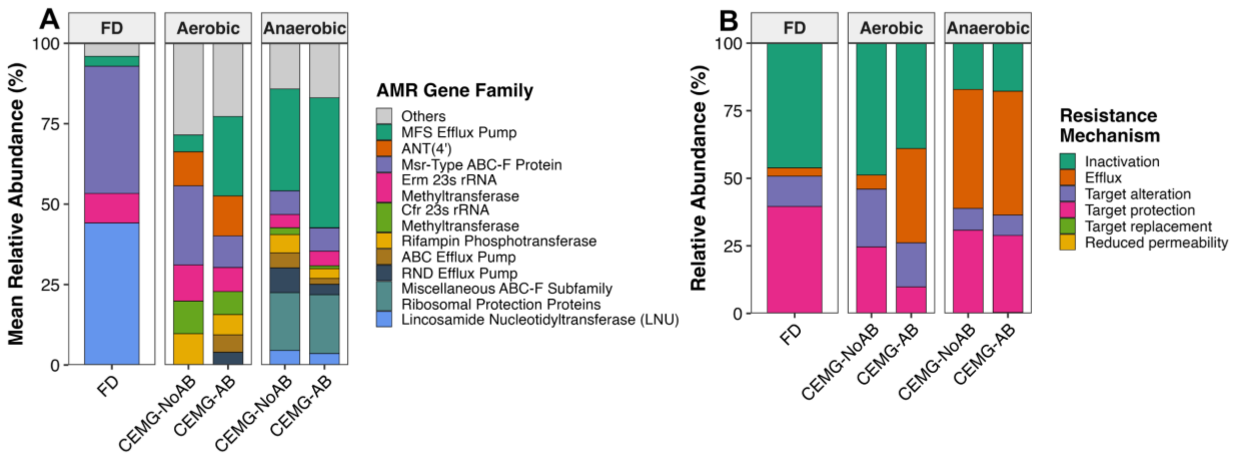


**Fig S5 AMR gene family composition and resistance mechanisms across enrichment conditions. (A)** Mean relative abundance of ARG gene families across FD, CEMG-NoAB and CEMG-AB under both aerobic and anaerobic conditions, expressed as a percentage of total ARG-mapping reads within each group. Within each oxygen condition, bars represent CEMG-NoAB (left) and CEMG-AB (right). **(B)** Relative abundance of resistance mechanisms corresponding to the gene families shown in (A), using the same grouping structure.

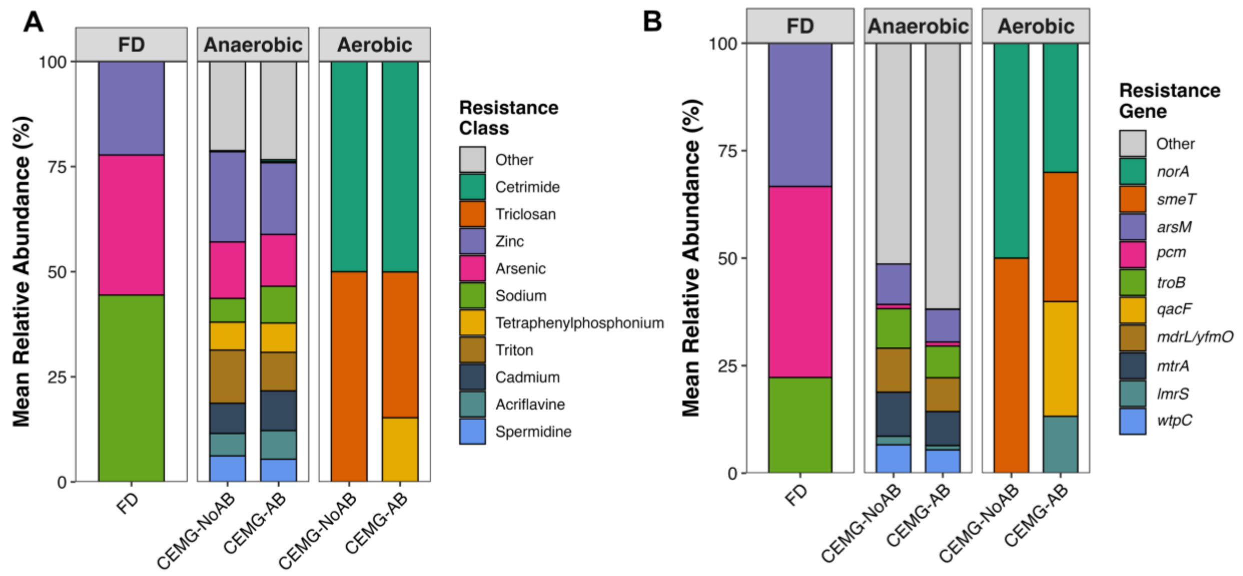


**Fig S6 Heavy metal resistance profiles associated with ARG-MGE-bearing contigs across enrichment conditions**. **(A)** Relative abundance of heavy metal resistance classes and **(B)** heavy metal resistance genes, across FD, CEMG-NoAB, and CEMG-AB under aerobic and anaerobic conditions.

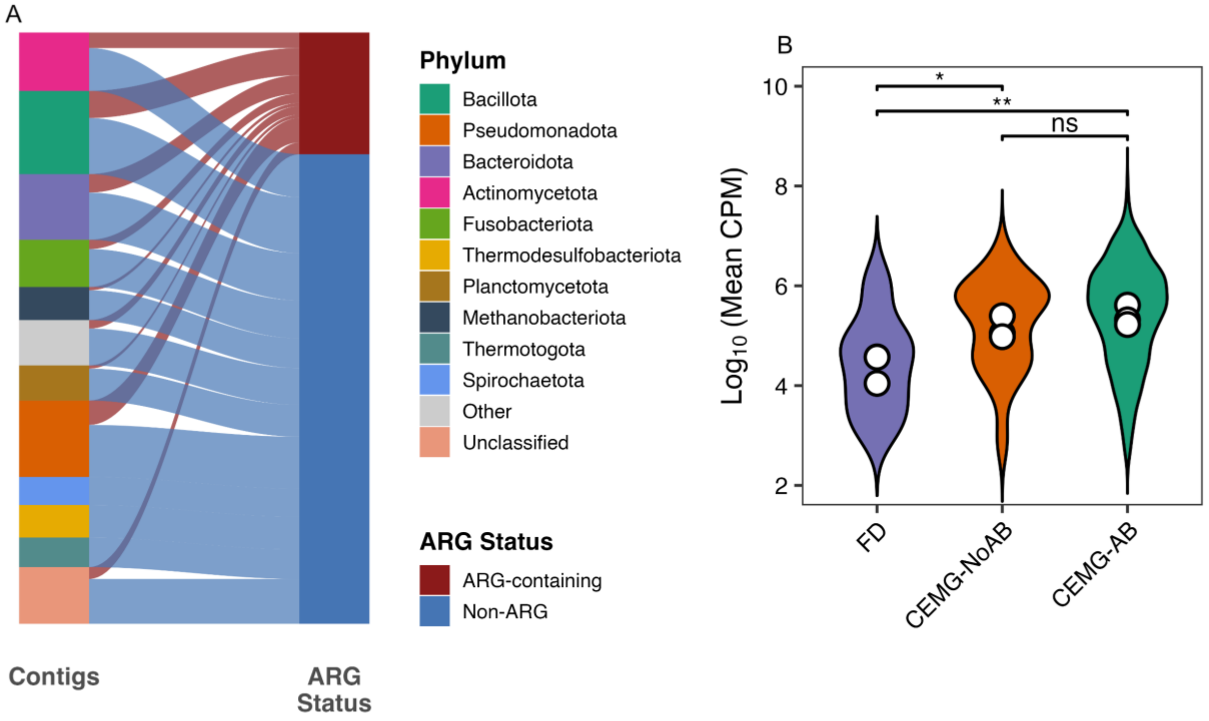


**Figure S7 Taxonomic distribution and abundance of ARG-bearing contigs across hybrid assemblies and enrichment conditions.** (**A**) Distribution of hybrid-assembled contigs by ARG status and assigned phylum. Contigs are partitioned into ARG-containing and non-ARG groups and linked to their corresponding taxonomic classifications. The non-ARG containing stratum was scaled down 20-fold for better visualization. **(B)** Abundance of ARG-MGE-associated contigs across enrichment conditions. $Log_{10}$-transformed mean count per million (CPM) is shown for FD, CEMG-NoAB, and CEMG-AB. Violin plots represent the distribution within each group, and white circles indicate group means. Statistical analysis was performed using a linear mixed-effects model. Pairwise group differences were evaluated using estimated marginal means (emmeans), with *p*-values corrected for multiple testing using the Benjamini-Hochberg procedure. Adjusted significance levels are denoted as ns (not significant), $p < 0.05$ (*), and $p < 0.01$ (**).

**Title: Optimizing a Culture-Enriched Hybrid Metagenomics Pipeline to Assess the AMR Footprint of Livestock Manure in Anaerobic Digestate**

*Nahidur Rahman*[1#], *A S M Zisanur Rahman*[1#], *David B. Levin*[2], *Tim A. McAllister*[1,3], *Nazim Cicek*[2], and *Hooman Derakhshani*[1*]

[1]Department of Animal Science, University of Manitoba, Winnipeg, MB, Canada
[2]Department of Biosystems Engineering, University of Manitoba, Winnipeg, MB, Canada
[3]Lethbridge Research Centre, Agriculture and Agri-Food Canada, Lethbridge, AB, Canada

### Supplementary Tables

**Table S1. Sample metadata and co-assembly group assignments.** Barcode, sample descriptions, abbreviations, antibiotic treatment, co-assembly group designation, sample name, and biological replicate number for all 81 samples used in this study (FD, n = 3; CEMG-NoAB, n = 6; CEMG-AB, n = 72).

Available on: https://doi.org/10.6084/m9.figshare.32942219

**Table S2. Illumina sequencing depth summary by sample group.** Mean and standard deviation of read pairs per sample for each group (FD, CEMG-NoAB, CEMG-AB) following quality control.

| Group | n | mean_pairs | sd_pairs |
|---|---|---|---|
| CEMG-NoAB | 6 | 9613166.33 | 4668001.91 |
| CEMG-AB | 72 | 11313766.9 | 3729106.53 |
| FD | 3 | 31919217.3 | 1200425.42 |

**Table S3. Taxonomic relative abundance profiles across all samples.** Clade-level relative abundance values (%) for all 81 samples generated by MetaPhlAn v4.2.2 against the CHOCOPhlAnSGB vJan25 marker gene database. Values represent the estimated relative abundance of each taxonomic clade as a percentage of the total classified community within each sample, as calculated by MetaPhlAn from clade-specific marker gene profiles; a value of 0 indicates the clade was not detected in that sample. Rows represent taxonomic clades spanning phylum to species level; columns represent individual samples.

Available on: https://doi.org/10.6084/m9.figshare.32942243

**Table S4. Dose-response model summary for taxonomic richness.** Best-fit model (linear or quadratic), model coefficients, p-values, $R^2$ values, and AIC difference (ΔAIC) for the association between antibiotic concentration (1X, 10X, 100X) and genus-level Shannon diversity, fit separately for each of the six antibiotic treatments. Model selection was based on ΔAIC, with linear models preferred when ΔAIC < 2.

Available on: https://doi.org/10.6084/m9.figshare.32942258

**Table S5. Antibiotic-enrichment-specific species detected exclusively in CEMG-AB.** Species detected in at least two CEMG-AB samples (relative abundance > 0) and absent from all FD and CEMG-NoAB samples (n = 91). Columns: species name, number of CEMG-AB samples in which the species was detected (n_AB), and maximum relative abundance across CEMG-AB samples (max_pct, %).

Available on: https://doi.org/10.6084/m9.figshare.32942270

**Table S6. ARG abundance across sample groups quantified by CARD-based read mapping.** Summary of sequencing depth, CARD-mapped read counts, and ARG abundance (expressed as counts per million total reads, CPM) across FD, CEMG-NoAB, and CEMG-AB. CPM was calculated as the number of reads mapping to CARD reference sequences divided by total library reads, multiplied by $10^6$.

| Group | n | Mean total reads (x$10^6$) | Mean CARD-mapped reads | Mean CPM | SD CPM | Median CPM | IQR CPM | % reads mapping to ARGs (mean) |
|---|---|---|---|---|---|---|---|---|
| FD | 3 | 63.8 | 974 | 15.4 | 7.3 | 14.1 | 7.2 | 0.15% |
| CEMG-NoAB | 6 | 19.2 | 2,365 | 124 | 86.4 | 112 | 36.7 | 1.03% |
| CEMG-AB | 72 | 22.6 | 3,644 | 160.4 | 94.8 | 139.1 | 128.4 | 1.58% |

**Table S7. Contig-level ARG-MGE co-localization associations across hybrid assemblies.** Each row represents a single association between an ARG, and an MGE feature detected on the same contig. Columns describe the contig of origin, assembly unit, biological replicate, sample group, ARG symbol, ARG drug class, ARG genomic coordinates (arg_start, arg_stop), consensus detection flag (consensus_hit; TRUE = detected by both RGI and AMRFinderPlus), RGI confidence cut-off (cut_off), alignment method (method), MGE class, detection tool, MGE feature coordinates where available (mge_start, mge_stop), distance in base pairs between the ARG locus and the MGE feature (distance_bp; 0 = overlapping), association window applied (2 kb for insertion sequences and recombinases; 25 kb for ICEs and IMEs; contig-level for plasmid and phage classifications), and annotation detail (detail; includes IS family, geNomad score, taxonomic lineage for viruses, or HMM profile hits and e-values for signature-gene classifications). NA values in coordinate and distance fields indicate contig-level classifications for which sub-contig coordinates were not resolved.

Available on: https://doi.org/10.6084/m9.figshare.32942300

**Table S8. ARG-MGE co-localization patterns stratified by drug class, gene symbol, bacterial genus, and MGE class**. Each row represents a unique combination of resistance drug class, ARG gene symbol, bacterial genus, and MGE class, with the corresponding number of ARG-bearing contig instances observed across all replicates and sample groups. Genus assignments are based on Kraken2 taxonomic classification of ARG-bearing contigs against the PlusPFP database (build 20250402). MGE classes were assigned using the multi-tool pipeline described in the Methods.

Available on: https://doi.org/10.6084/m9.figshare.33007574
